# Plate design confounds donor variation and model evaluation in single-cell perturbation screens

**DOI:** 10.64898/2026.11.04.764088

**Authors:** Ghazal Ghajari, Krishnaprasad Thirunarayan, Fathi Amsaad

**Affiliations:** Department of Computer Science and Engineering, Wright State University, Dayton, OH 45435, USA

**Keywords:** single-cell perturbation, inter-individual variation, well effects, negative controls, benchmarking, cytokines, peripheral blood mononuclear cells, experimental design

## Abstract

Single-cell screens of immune cells from many donors are used to study why individuals respond differently and to benchmark response models. Each donor–perturbation combination typically occupies one well, controls are grouped, and donors share one layout, so technical effects can look biological. Using a 90-cytokine screen, two compound screens, four cohorts and simulations, we show that well effects explain most replicate-control variation in abundant cell types, controls carry a shared offset, and identical control wells differ by hundreds of differentially expressed genes at a standard threshold, about as many as two receptor-negative proteins do. Negative controls, design-aware estimators and perturbation-specific evaluation change the interpretation of four recent analyses. A deep-learning model that scores below a perturbation-agnostic predictor under standard evaluation outperforms a simple baseline on perturbation-specific differences in two of three cell types (significantly in one with stricter controls), but predicts less well the component shared with inert wells.

## INTRODUCTION

Single-cell perturbation screens have expanded from cell lines to primary immune cells collected from many individuals. A recent cytokine screen profiled nearly ten million peripheral blood mononuclear cells (PBMCs) from 12 donors exposed to 90 cytokines,^1,2^ and a compound screen of PBMCs from three donors now serves as a community benchmark. ^3^ Such data are used both to ask how individuals differ in their response to the same stimulus ^2,4^ and to train and compare models that predict the response of an unseen donor, cell type or perturbation.^5–9^ These uses matter beyond the screens themselves: differences between donors are a starting point for understanding why individuals respond differently to the same immune signal or drug, and response models are being developed to prioritize perturbations before they are tested. Technical effects that look biological can mislead both.

This work began with a question about recent decomposition methods, which separate a perturbation response into a program shared across perturbations and a component specific to each: ^10,11^ which component carries the differences between individuals? To answer it, we first had to establish whether the measured differences between donors were genuine. In several cases they were not, and this paper is about why.

The cause is the layout of these screens. Batch effects are well recognized in genomics,^12^ and in single-cell studies they confound comparisons whenever plates or batches coincide with the groups being compared. ^13,14^ In high-throughput screening, positional effects within plates are known to bias comparisons, particularly when all controls sit together at the plate edge. To avoid this, replicate wells and distributed or randomized control positions are recommended;^15,16^ in image-based profiling, well position is likewise balanced by placing replicates in different positions. ^17^ In probe-based single-cell assays, sample barcodes have recently been shown to leave a reproducible signature of their own, ^18^ but whether this holds for other barcoding schemes is unknown. Plate-based perturbation screens add three features that have received little attention: each donor–perturbation combination occupies a single well shared by all cell types in the sample; control wells are placed together in one region of the plate; and the layout is identical for every donor, so a perturbation always occupies the same well and carries the same barcode. Together, these features let a technical signature recur in every donor and pass tests that treat donors as replicates.

Here we quantify how these features, together with common analysis choices, create technical signals that look biological: spurious similarity, when a well’s signature recurs across the cell types of a well or across donors, and spurious differences between donors (Table 1). We examine these effects in two further plate-based datasets and, for donor-level effects, in four independent cohorts, and test a corrected pipeline and a randomized layout on simulated data with known ground truth. We then re-examine four publicly reported analyses, namely a coordination analysis of donor responses, ^4^ a benchmark of compound responses,^3^ the DEG counts of the Cytokine Dictionary ^2^ and the evaluation of the State model ^5^ (with the simple method Rhaister ^6^ rescored alongside), and show that the interpretation of each of these four analyses changes, partly or fully, once the design is taken into account. Finally, we examine a baseline interferon effect that persisted after technical corrections within one dataset but did not replicate, and give practical recommendations for future screens. The principle that treatments must not be confounded with position is long established; our contribution is to quantify its consequences in single-cell perturbation screens, to provide negative controls that isolate them, namely receptor-negative proteins and identical control wells, and to show how they change published analyses.

**Table 1.** Design features, technical effects, affected analyses and remedies.

| Feature | Technical effect (this study) | Analyses affected | Remedy |
|---|---|---|---|
| One well per donor and perturbation, shared by all cell types | Well noise: 32% to 83% (Parse), 17% to 69% (OP3) and a median of 38% (Tahoe-100M) of replicate-control differences, correlated across cell types or cell lines (Figures 2 and S6) | Coordination across cell types within donors; split-half ceilings (Figures 4A, 4B and 5A) | Replicate wells; baselines from wells without an active protein; ceilings from separate wells |
| Controls grouped in one row (Parse; also Tahoe-100M) | Offset shared by every comparison with the control, beyond the noise of the control mean; differences growing along the row (Figures 1 and S5B) | Responsiveness calls and correlation-based benchmark scores against PBS (Figure 5) | Controls distributed across the plate; inert-protein reference; removal of the shared component before scoring |
| Identical layout for every donor | Well-specific signatures that recur in every donor or on every plate, from position, barcode or reagent (Figures 4E, S5 and S6) | DEG counts; compound responses in OP3; any test with donors as replicates (Figures 4C to 4E) | Positions randomized or assigned to donors; receptor-negative protein controls; DEG counts read against the floor |
| Analysis choices: estimator, unequal cell numbers, processing batch | Spurious donor differences in response size and shape (Figure 3) | Associations between baseline interferon and response shape (Figure 6) | Cross-half estimators, equal cell numbers, within-batch comparisons |

## RESULTS

### In current screens each perturbation is tied to one well, one barcode and one position

The Parse cytokine screen comprises 9,697,974 PBMCs from 12 healthy donors treated for 24 h with one of 90 cytokines or PBS.^1,2^ Each donor was profiled on a single 96-well plate, every cytokine occupies the same well on every plate, and PBS fills six wells at the plate edge, H7 to H12 (Figure 1A). The OP3 screen, which underlies the Open Problems benchmark for chemical perturbations, ^3^ follows the same principle: PBMCs from three donors were distributed over six plates and treated with 144 compounds, three of which are controls occupying columns 1 to 3, and each of the other 141 compounds occupies the same well on every plate on which it appears. ^3^ In both designs, a perturbation is inseparable from its well and plate position and, in Parse, from its first-round barcode, because split-pool barcoding labels the cells of each well in the first round. ^19^ Throughout, we call the part of every response that inactive wells also show the shared component, estimated from the assumed-inert proteins; the response common to all perturbations contains this component and, in addition, biology shared among active perturbations.

**Figure 1.**
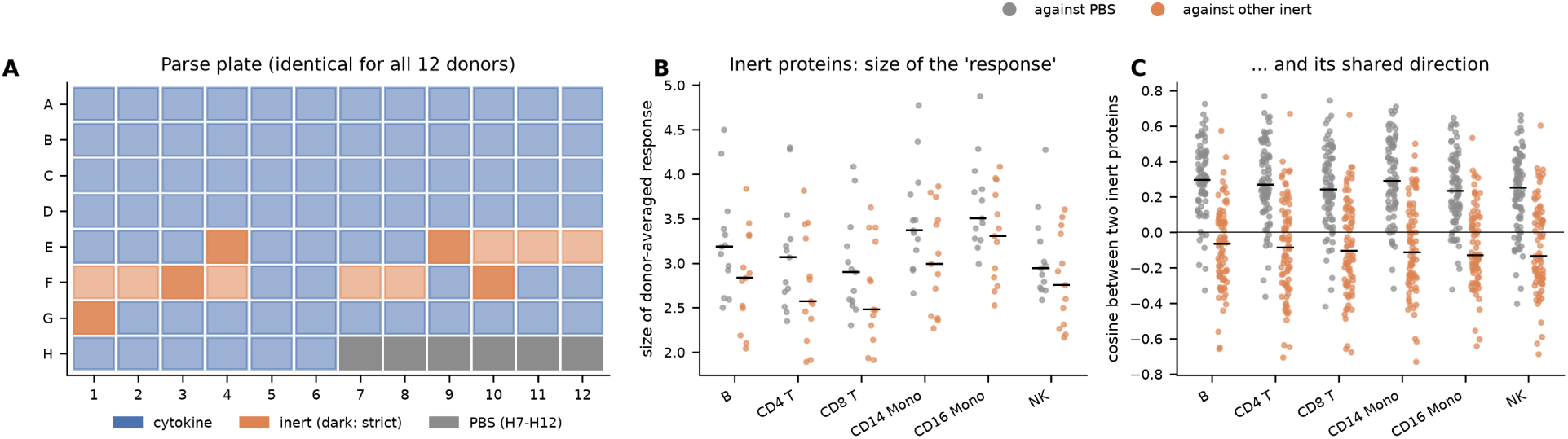
PBS wells carry an offset beyond the noise of their own mean. (A) Plate layout of the Parse screen, identical for all 12 donors. Each cytokine occupies one well; the 13 assumed-inert proteins (dark orange, the five receptor-negative proteins) and the six PBS wells in row H are highlighted. (B) Size of the donor-averaged response of each inert protein against PBS (grey) and against the mean of the other inert proteins (orange), per cell type, on the 2,000 genes most expressed in PBS cells. One point per protein; lines, medians. (C) Cosine between the responses of pairs of inert proteins, computed against PBS (grey) and against the other inert proteins (orange). One point per pair; lines, medians. See also Figure S1.

### PBS wells carry a shared offset, and most of the apparent response to an inert protein is not shared with other proteins

With PBS as the reference, nearly all 90 cytokines appeared responsive in nearly every cell type, which is biologically implausible. To find the cause, we examined 13 proteins that are not expected to act on PBMCs, including Noggin, EPO and TPO, and asked how large their apparent responses were and whether these were shared across proteins.

Against PBS, the responses to different inert proteins were partly aligned (median pairwise cosine 0.28, range 0.23 to 0.30 across cell types; Figure 1C), twice the value of about 0.14 (1/7) expected from the noise of the six PBS wells alone if all wells were equally noisy. Against the other inert proteins, this shared direction disappeared (median cosine *−*0.11, close to the value of approximately *−*1*/*12 expected by construction), so PBS wells differ from the rest of the plate by a common offset. The response to an inert protein was 12% larger against PBS than against the other inert proteins (median size 3.19 versus 2.84; Figure 1B), whereas noise of the six PBS wells alone would make it only about 4% larger; because sizes add in quadrature, the remainder corresponds to a shared component of about 40% of the size of an inert response, if all wells are equally noisy. The offset may arise from the grouping of the PBS wells in one row, as expected for controls placed together at the plate edge, ^15,16^ from PBS wells being noisier than other wells, for example at the edge, or from a reagent component that protein wells received but PBS wells did not, such as a carrier protein, buffer or trace endotoxin; the design does not separate these explanations. Endotoxin is unlikely to be the main source: LPS-responsive genes (20 genes, for example IL1B, TNF and CXCL8; STAR Methods) were lower, not higher, in inert-protein wells than in PBS wells, most clearly in monocytes (*z* = *−*3.8 to *−*4.4; STAR Methods). Whatever its source, this component is shared by every comparison with PBS and does not reflect a response to any particular protein. Most of the apparent response to an inert protein was instead specific to its own well (see below).

The two observations have distinct consequences. For identifying responsive cytokines, the per-well signature is the main obstacle, so we derived the response threshold from differences between inert proteins rather than from a significance test against a control. For evaluating prediction models, the offset matters more than its size suggests, because it is identical for every cytokine and correlation-based benchmark metrics are insensitive to the size of a shared component.

To assess whether all 13 proteins are plausibly inert, we examined the expression of their receptors in PBS-treated cells (Figure S1; STAR Methods). The receptors of five proteins were expressed in at least one main cell type, the status of three was uncertain, and the ligand-binding receptors of TPO, PSPN, EPO and EGF were essentially absent, while Megalin is itself a receptor rather than a signaling ligand. We therefore defined TPO, PSPN, EPO, EGF and Megalin, the receptor-negative proteins, as a strict inert set, repeated every analysis with these five proteins alone, and report both versions for the main analyses.

With the mean of the inert proteins as the reference, we called a cytokine responsive only when its difference from this reference exceeded the differences observed between inert proteins. This gave biologically plausible results: 8 to 19 responsive cytokines per main cell type, with interferons active in almost all cell types and the expected responses to B cell, monocyte and common gamma-chain cytokines.

### Well-to-well noise accounts for most of the difference between replicate control wells

Because each cytokine occupies only one well per donor, well noise cannot be measured from cytokine conditions, but the replicate PBS wells allow it. Halves of cells from different PBS wells differed more than halves from the same well: across the main cell types, the well accounted for 32% to 83% of the difference between replicate wells on the 2,000 genes most expressed in PBS cells (15% to 74% on the highly variable genes used for donor comparisons), and for 79% and 83% in the two most abundant cell types, CD4 T cells and CD14 monocytes (Figure 2A). OP3, which has eight DMSO wells per plate, gave similar estimates: 67% to 69% in T and myeloid cells, 32% in B cells and 17% in NK cells. In a pooled design, the corresponding technical unit contributed much less: in the cohort of Aquino et al., ^20^ in which each sample was stimulated separately, pooled with others and processed on two libraries, the library accounted for only 5% to 10% of the difference between the two replicate libraries of unstimulated samples of 222 donors (Figure S2A), although this comparison does not capture noise that arises before pooling, such as that of the stimulation well.

**Figure 2.**
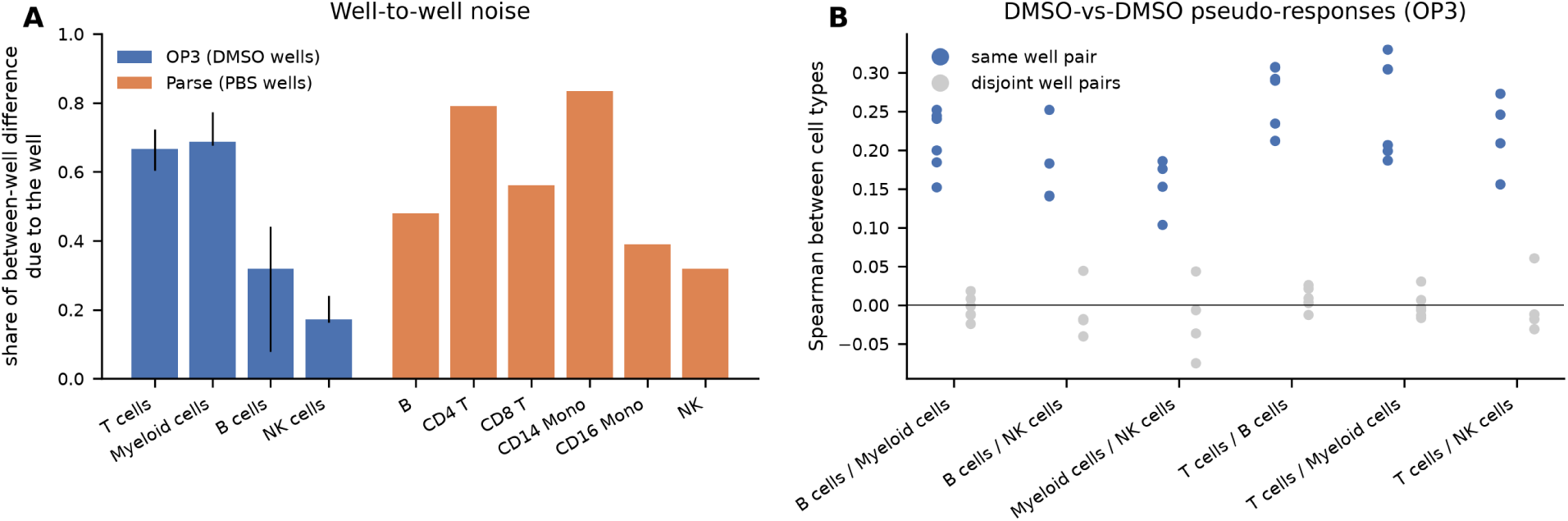
Well-to-well noise accounts for most of the difference between replicate control wells. (A) Share of the difference between replicate control wells that is due to the well itself, (between *−* within)/between, for OP3 DMSO wells (bars, median over six plates; lines, range) and Parse PBS wells (median over 12 donors), on the 2,000 most highly expressed genes (in all cells for OP3 and in PBS cells for Parse). (B) OP3: Spearman correlation between cell types of the difference between two DMSO wells, for cell types from the same pair of wells (blue) and from disjoint pairs (grey). One point per plate; pairs involving NK cells include only plates with enough NK cells.

Because all cell types in a sample share a well, well effects should be correlated across cell types. In OP3, for a given pair of DMSO wells, so that no treatment was involved, the difference in one cell type correlated with that in another (median Spearman correlation 0.21), whereas disjoint pairs of wells were uncorrelated (*−*0.01), on every plate and for every pair of cell types (Figure 2B).

The two OP3 positive-control compounds, belinostat and dabrafenib, also occupy all eight rows of every plate, which allows the same estimate for active treatments. Their median well energy (the part of the squared difference between two wells that exceeds sampling noise; STAR Methods) was 0.20 to 0.81 times that of DMSO on the same plates and in the same cell types, although dabrafenib exceeded DMSO on single plates (Figure S3A). Well noise therefore did not increase with the response on average, and estimates from replicate control wells, as used in our corrected pipeline, do not on average underestimate the well noise of active treatments.

### Estimator choice, cell numbers and processing batch each create donor differences

Four Parse donors have high baseline expression of interferon-stimulated genes, a difference examined in the last section of the Results, so we use a baseline interferon score (Method details) to illustrate how analysis choices create donor differences. Whether two donors differ in the size or the shape of their response follows from the identity ||*a – b*|| ^2^ = (*|a| −|b|*)^2^ + 2*|a||b|*(1 *−* cos(*a, b*)), which partitions their difference into a size and a shape component without assuming a model. Sampling noise contributes to both components, and the way it is removed affects the result.

A simple asymmetric estimator, which compares one half of a donor’s cells with the nearly noise-free mean of the other donors (STAR Methods), produced in Parse impossible negative shape values in up to 31% of cases and inflated a group contrast by a factor of 1.2 to 4. In 1M-scBloodNL, a cohort of 120 donors exposed to three pathogens,^21^ it produced correlations of 0.3 to 0.5 between the baseline interferon score and shape deviation, which fell to approximately zero when the estimator was made symmetric and donors were compared only with others processed on the same day. In the same cohort, the interferon score varied strongly between processing dates, and in 90 donors from Randolph et al. ^22^ it varied far more between batches than between ancestry groups (batch means *−*0.49 to 0.70, versus *−*0.04 and 0.04 for the two ancestry groups; Figure 3A). In 222 donors from Aquino et al., ^20^ processing run explained 16% of its variation (permutation *p* = 0.002), ancestry 3% (*p* = 0.033), and post-thaw cell mortality, cytomegalovirus status, age and sex less than 1% each (Figure S2B).

**Figure 3.**
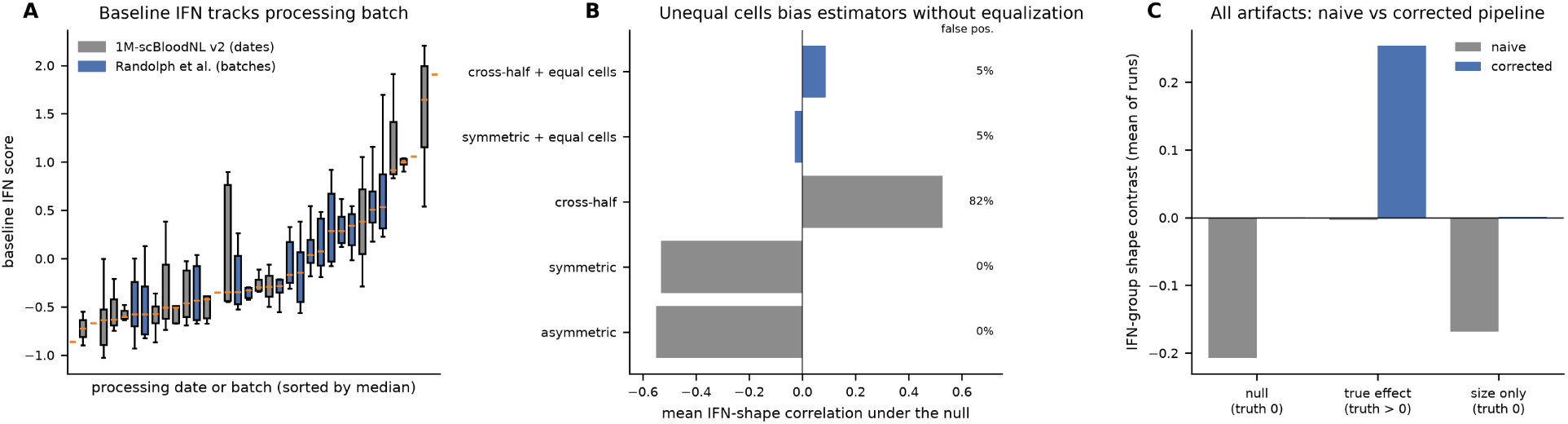
Estimator choice, cell numbers and processing batch each create donor differences. (A) Baseline interferon score of donors grouped by processing date in 1M-scBloodNL v2 (grey) and by batch in Randolph et al. (blue). Boxes, interquartile range; lines, medians. (B) Simulation without any true effect in which high-interferon donors have fewer cells: mean correlation between the interferon score and shape deviation for five estimators, and the share of runs with a significant positive association (right). 40 simulated datasets of 24 donors per setting. (C) Simulation with all artifacts present: mean interferon-group shape contrast for the naive and corrected pipelines when there is no effect, a true shape effect, or differences in size only. 40 simulated datasets per setting.

We next simulated 24 donors with known ground truth, with 40 datasets per setting (Table S1). When high-interferon donors had fewer cells, every estimator without equalization of cell numbers was biased, in a direction that depended on the estimator: the asymmetric and symmetric estimators biased shape deviation downward, whereas an estimator built only from independent halves biased it upward and produced false positives in 82% of runs (Figure 3B). A processing-date effect produced false positives in 50% of runs. Well noise made donors that differed only in response size appear to differ in shape, so that only 30% of the true size share was recovered. With all artifacts present, the naive pipeline produced a group difference where none existed and detected a real one in only 10% of runs; the spurious difference was negative, and so not counted as a false positive by our one-sided criterion, because the naive estimator biases shape deviation downward when donors have fewer cells (Figure 3B). The corrected pipeline (inert reference, cross-half estimator, equal cell numbers, within-batch comparisons and a well-noise correction from replicate control wells) kept false positives, defined as significant positive associations, at 3% and detected the real effect in 65% of runs (rates over 40 runs; Figure 3C shows the corresponding group contrasts).

A second simulation asked which of these artifacts a different layout would remove; it illustrates the expected effect of randomization rather than testing it. Each plate position carried a signature that was identical on every donor plate. With a fixed layout, as in Parse, the responses to inert proteins agreed between two groups of 12 donors (mean correlation 0.87), two groups of identical control wells differed by 209 DEGs of 500 genes on average, and a predictor that ignored the cytokine reached a correlation of 0.21 with the observed responses; with its own random layout for each donor plate, these values fell to 0.01, 0.05 and 0.01 (Figure S4A–C). The coordination of inert responses across the cell types of one donor persisted, because the cell types of a well share its noise wherever the well lies: it was 0.89 in both layouts, but its donor-shuffled null was 0.31 with the fixed layout, which retained the position signature common to all donors, and *−*0.01 with the randomized layout, giving excesses over the null of 0.56 and 0.89 (Figure S4D). A fixed layout therefore inflates the null as well as the observed statistic, and a donor-shuffled test cannot show how much of the coordination is technical. Randomization removes the artifacts that recur across donors, but it does not replace replicate wells and estimators built from independent halves, nor does it remove the common component rewarded by benchmark scores: in the simulation, the predictor that ignored the cytokine was estimated from other donors, whereas in benchmarks it is the query context’s own mean response, which contains biology shared among perturbations and the noise of that context’s own controls. Removing the shared component in the evaluation therefore remains necessary.

### Inert proteins and identical control wells reproduce most of a reported coordination signal

Zahid ^4^ reported that, within a donor, responses to every one of the 90 cytokines are coordinated across cell types beyond what is expected when donor labels are shuffled. Shuffling donor labels, however, pairs cell types from different wells and so removes any effect shared by the cell types of a single well; the test therefore cannot distinguish shared well effects from donor-level organization.

We reproduced the analysis: as reported, every cytokine exceeded the shuffled null, and so did every inert protein (Figure 4A), with an excess of 70% to 80% of that of immune cytokines in gene space, regardless of the reference and of the inert set. A comparison without any stimulus, between two groups of PBS wells (H7 to H9 versus H10 to H12), gave an excess of 0.105 (*p ≤* 0.002, the smallest value attainable with 500 permutations), about 90% of the median excess of immune cytokines (Figure 4A) and 60% to 90% of it across five comparison spaces (Figure 4B); the cytokines with the strongest transcriptional effects, such as interferon-*β*, IL-2 and IL-15, fell at this technical level. With all 13 inert proteins as controls, immune cytokines lay modestly but significantly above them (Mann-Whitney *p* = 0.004), so a smaller component above the technical level does exist, although it is not concentrated in the cytokines with the strongest effects; with the five receptor-negative proteins the difference was not statistically detectable (*p* = 0.057), which, given the low power of a test with five controls, we interpret as a small difference rather than an absence of difference. The result did not depend on the comparison space: in gene space, in two PCA spaces learned on Parse, in a random 150-dimensional space and in a 150-component space learned, as in the original analysis, from baseline donor profiles of an independent cohort (222 donors from Aquino et al.; ^20^ 3,424 shared genes), inert proteins reached 74% to 95% of the immune excess across spaces, references and inert sets (Figure 4B). In this external space, immune cytokines did not lie significantly above the inert proteins (Mann-Whitney *p* = 0.09 with the PBS reference and 0.14 with the inert reference).

**Figure 4.**
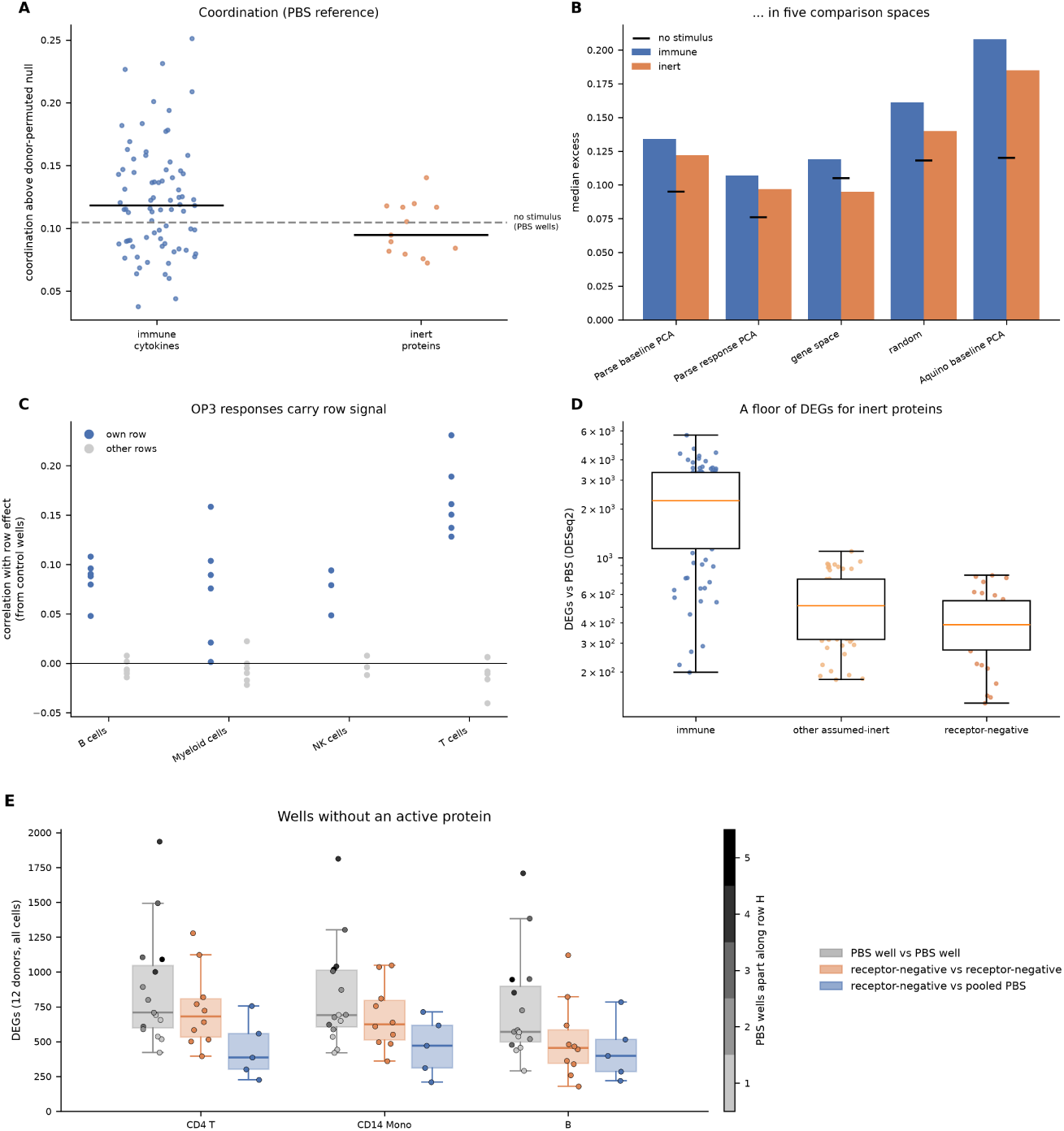
Shared wells and fixed layouts contribute to three published signals and create a floor of DEGs without any protein. (A) Coordination across cell types within donors, above the donor-shuffled null, for 77 immune cytokines and 13 inert proteins (PBS reference). Lines, medians; dashed line, the same statistic for a comparison between two groups of PBS wells without any stimulus. 500 permutations. (B) Median excess for immune cytokines and inert proteins (PBS reference, all 13 inert proteins) in five comparison spaces, including a 150-component space learned on baseline profiles of an independent cohort (Aquino et al.); black marks, the stimulus-free comparison. (C) OP3: correlation of compound responses with the row effect of their own row (blue) and of other rows (grey), both estimated from control wells. One point per plate; row permutation test with 1,000 permutations. (D) DEGs against PBS (pseudobulk DESeq2, adjusted *p <* 0.05, | log_2_ fold change*| >* 0.25) for 13 well-characterized immune cytokines (IFN-*β*, IFN-*γ*, IL-2, IL-4, IL-7, IL-10, IL-15, IL-21, GM-CSF, M-CSF, IL-1*β*, TNF-*α* and CD40L), the eight other assumed-inert proteins, with expressed or uncertain receptors (HGF, PRL, VEGF, IGF-1, Decorin, Noggin, GDNF and FGF-*β*) and the five receptor-negative proteins, in six cell types; all 13 inert proteins taken together had a median of 415 DEGs. Boxes, interquartile range. (E) DEGs between two wells without an active protein, with all 12 donors and all cells: pairs of PBS wells (grey; shade, distance along row H), pairs of receptor-negative proteins (orange) and receptor-negative proteins against pooled PBS (blue). Genes with at least 10 counts per million in PBS cells. Most DEGs have small effects; counts with stricter fold-change thresholds are given in the text. See also Figure S5.

### Compound responses in a benchmark screen carry the signal of their plate row

The OP3 benchmark, published in the NeurIPS 2024 Datasets and Benchmarks track, asks models to predict differential expression statistics for each compound relative to DMSO, computed with limma with plate but no term for row or column in the design (STAR Methods). ^3^ Because the three control compounds occupy every row, row effects can be estimated from control wells alone; they were weak but reproducible across plates. Compound responses correlated with the row effect of their own row, estimated from entirely separate wells, at a median of 0.08 to 0.16 across cell types (up to 0.23 on single plates), and with the effects of other rows at a median of *−*0.01 (row permutation *p <* 0.05 in 19 of 21 combinations of plate and cell type, comprising T, B and myeloid cells on six plates and NK cells on the three plates with sufficient cells; Figure 4C). The published targets carried the same signal: their log fold changes correlated with the row effect of the compound’s own row at a median of 0.04 to 0.15 across cell types, compared with 0.005 to 0.013 for other rows (row permutation *p ≤* 0.0005 in every cell type). In the benchmark’s own metric this signal was small. A predictor that used only position, the mean target of the other compounds in the same row, scored 0.009 to 0.028 higher (mean row-wise Pearson correlation) than the same predictor built from random groups of compounds of equal size (*p* = 0.001 to 0.016), against scores of about 0.20 to 0.31, and scored below the mean of all other compounds, which averages over more compounds. Because each compound occupies the same well on every plate, this component cannot be separated from the compound itself.

### Wells without an active protein yield hundreds of DEGs

The Cytokine Dictionary, the study that released the Parse screen, summarizes each combination of cytokine and cell type by the number of differentially expressed genes (DEGs) relative to PBS.^2^ Counting DEGs in the same way, with pseudobulk DESeq2^23,24^ and the thresholds of the original study, the 13 well-characterized immune cytokines of Figure 4D, chosen before the analysis, yielded a median of 2,241 DEGs, all 13 inert proteins taken together a median of 415 and the five receptor-negative proteins a median of 389, with DEGs in every cell type (Figure 4D); with the other inert proteins as the reference, 376 remained. DEG counts therefore distinguish strong cytokines from inert proteins, but every well has a floor of several hundred DEGs that no choice of reference removes.

This floor does not require a protein. Pairs of PBS wells, which received an identical treatment and differ only in position, first-round barcode and handling, differed by a median of 690 DEGs (all 12 donors and all cells, in B cells, CD14 monocytes and CD4 T cells), compared with 597 for two receptor-negative proteins and 399 for a receptor-negative protein against pooled PBS (Figure 4E); even adjacent PBS wells differed by 456 to 536 DEGs, about as much as two receptor-negative proteins (455 to 681). Most of these DEGs had small effects (median | log_2_ fold change| 0.33 to 0.37), close to the threshold of the original study. With a threshold of 0.5, two PBS wells still differed by a median of 112 DEGs and two receptor-negative proteins by 90; with a threshold of 1, by 12 and 10. Five cytokines chosen in advance as canonical strong responders (IFN-*β*, IFN-*γ*, IL-2, IL-15 and IL-4) yielded medians of 3,355, 1,346 and 349 DEGs against pooled PBS at the three thresholds, and a receptor-negative protein 399, 50 and 7, so the floor amounts to about 12% of a strong response at the original threshold but to only about 2% at a threshold of 1. A stricter threshold therefore shrinks this floor, whereas the coordination across cell types, the shared component rewarded by correlation-based scores and the inflated split-half ceilings do not involve DEG counts. Along row H, the number of DEGs between PBS wells increased approximately twofold from adjacent to the most distant pairs, although not monotonically: counts peaked at a distance of four wells, and the largest distance comprises a single pair (Mantel *r* = 0.69 to 0.73, *p* = 0.019 to 0.025; Figure S5B). The association persisted when any single well was removed (*r* = 0.58 to 0.95), indicating a gradient along the row rather than one outlying well; because position and handling order coincide along the row, the two cannot be distinguished. For the receptor-negative proteins, which occupy mostly interior wells, DEG counts were unrelated to plate distance (Mantel *r* = *−*0.35 to *−*0.44, *p ≥* 0.92).

The responses to inert proteins were reproducible between two independent groups of six donors (correlation 0.37 to 0.79 for the same protein, *−*0.03 to *−*0.07 for different proteins) but unrelated to distance on the plate (Mantel *p* = 0.33 to 0.49), so in these mostly interior wells the signature is specific to each well rather than part of a smooth spatial gradient. It could arise from the well’s barcode, by analogy with the barcode effects described for probe-based assays, ^18^ from the reagent or from a weak true effect; the design cannot distinguish among these. Whatever its origin, because the layout is identical for every donor, the signature recurs across donors and passes any test that treats donors as replicates.

DEG counts also depend strongly on statistical power. With 100 cells per well, the same comparisons yielded 37 to 67 DEGs with all 12 donors and 10 to 14 with six donors (Figure S5A), so DEG counts cannot be compared between datasets, or between cell types, that differ in the number of donors or cells. Power explains most of the contrast with OP3, in which two wells of the same treatment yielded almost no DEGs (median 0 to 1 in B, T and myeloid cells; Figure S5C), whereas belinostat yielded 712 to 2,627 DEGs against DMSO in the same row. At a power comparable to that of OP3 (six replicates, matching its six plates, and about 100 cells per well), Parse comparisons still yielded 10 to 14 DEGs, so a modest difference between the two screens remains.

### The well effects recur in a large compound screen of cell lines

To test whether these effects are specific to screens of primary immune cells, we examined Tahoe-100M, ^25^ in which about 50 cancer cell lines were pooled in every well of 14 plates of 96 wells and exposed to 379 compounds. Each plate contains two DMSO wells. The first-round barcode, which identifies the well of the barcoding plate, places them at the same two positions on 13 of the 14 plates (positions 95 and 96, the last two wells of row H if positions run row by row) and at positions 23 and 24 (B11 and B12) on plate 13. The cell lines of a well share its handling and so play the role of the cell types. The well accounted for a median of 38% (interquartile range 25% to 51%) of the difference between the two DMSO wells of a plate, over 625 combinations of plate and cell line with at least 100 cells in each half, and the share was above zero on every plate (Figure S6A); because the two DMSO wells are adjacent, whereas the Parse estimates pool all pairs of PBS wells, including distant ones, the two estimates are not directly comparable. With the 14 plates as replicates, the two DMSO wells differed by a median of 947 DEGs per cell line at the threshold of the original Parse analysis, 269 with a threshold of 0.5 and 58 with a threshold of 1 (45 cell lines; Figure S6B). Measured as well energy, this difference amounted to a median of 20% of the response to a compound in 112 randomly chosen compound wells and to 5% of the strongest tenth of responses, and no compound response was smaller than this floor (Figure S6D). The difference between the two DMSO wells correlated across the cell lines of a plate (median Spearman correlation 0.43), as across cell types in OP3, and more weakly between cell lines on different plates (0.24; Figure S6C): these are different wells, but on 13 of the 14 plates they occupy the same two positions, so part of the difference recurs on nearly every plate, as expected when the layout is fixed. Plates 1 to 3 carry the same compounds in the same wells at three doses, so we compared the row effects of each with those of plate 4, which carries different compounds. Row effects of two plates agreed for the same row (median correlation 0.11) but not for different rows (*−*0.02; row permutation *p <* 0.05 in 50 of 136 combinations of cell line and pair of plates). The agreement was strong between plate 4 and the plate at the same, lowest dose (0.27; *p <* 0.05 in 42 of 46 cell lines) and weak for the two plates at higher doses (0.12 and 0.06; *p <* 0.05 in 7 of 46 and 1 of 44 cell lines, close to the rate expected by chance). This is consistent with a fixed positional signal that stronger compound responses mask, but the evidence rests mainly on one pair of plates, which may also have shared processing, and because positions were taken from the first-round barcode, these rows are those of the barcoding plate, which need not coincide with the plate on which cells were treated. Well noise shared by everything in a well, a DEG floor between identical control wells and a signature tied to barcode position thus appear in a third screen, with a different cell source, a different kind of perturbation and many more plates.

### Removing the shared component changes how State compares with simple base-lines

Prediction benchmarks on Parse score models by the Pearson correlation between predicted and observed responses relative to PBS.^5,6^ Part of every such response is common to all cytokines, and a model can score highly simply by predicting it. This part has two sources: biology shared among active cytokines, and the shared component defined above, which wells without an active protein also show, including the PBS offset.

On the donor split used by Rhaister, a method that predicts responses from observed statistics, ^6^ a predictor that returns the same vector for every cytokine, namely the mean of the responses already observed for the held-out donor (the context mean), reached 59% to 64% of the score of the additive baseline (0.339 versus 0.572 in our pseudobulk evaluation, and 0.306 versus 0.481 when re-scoring Rhaister’s released evaluation). A predictor with no information about either the cytokine or the donor, the mean response to inert proteins in the training donors (“inert shift”), scored 0.11 and 0.03 in the two evaluations (Figure 5B). The additive baseline also exceeded the split-half ceiling often used as an upper bound (0.572 versus 0.546). This ceiling is inflated, because two halves of the same well share that well’s effect: for inert proteins, halves of one well correlated at 0.41, whereas different wells correlated at *−*0.03 (Figure 5A). In OP3, halves of one DMSO well correlated at a median of 0.15 to 0.40 across cell types and different DMSO wells at 0.02 to 0.06, whereas for the strongest active compound, belinostat, the two correlations differed by only 0.01 to 0.06 (Figure S3B). The split-half ceiling is therefore inflated mainly for weak responses, which include most cytokines in most cell types.

**Figure 5.**
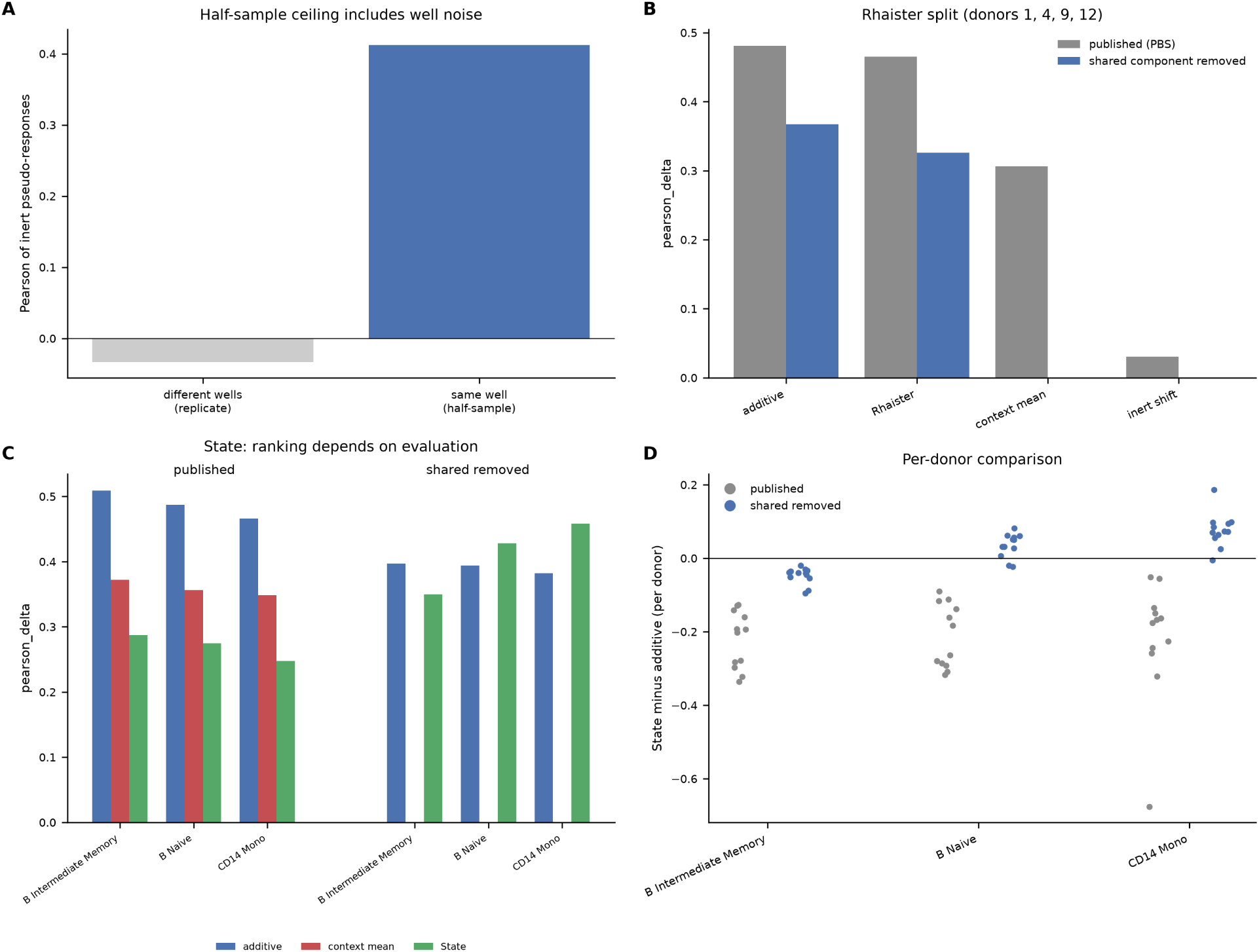
Removing the shared component changes how State compares with simple baselines. (A) Pearson correlation between inert-protein responses measured in two halves of the same well (the usual split-half ceiling) and in two different wells. (B) Donor split used by Rhaister, re-scoring Rhaister’s released evaluation: mean pearson delta over immune test cytokines for Rhaister, the additive baseline and two predictors without cytokine information, under the published evaluation (grey) and after removing the shared component (blue). After removal, predictors that return the same vector for every cytokine score zero by construction, so their bars are absent. (C) State’s published predictions on three held-out cell types, scored against one ground truth computed from raw Parse counts in State’s gene space: mean scores of the additive baseline, the cytokine-free context mean and State under both evaluations (the context mean scores zero after removal, as in B). (D) Difference between State and the additive baseline for each of the 12 donors, under both evaluations. Paired Wilcoxon tests on donor means.

We then rescored predictions after removing the shared component, subtracting the observed inert mean from the observed responses and each method’s own predicted inert mean from its predictions (STAR Methods); any prediction identical for every cytokine then scores zero. The corrected score measures how well a model predicts the difference between each cytokine and a cytokine without effect, which we consider the quantity response models aim to learn; because this is a choice, and because it ignores errors in the predicted shared component, we report both versions and a sensitivity analysis. Because the subtracted vector comes only from proteins without a detectable effect, biology shared among active cytokines is retained. This distinguishes the correction from Systema, which addresses systematic variation in genetic perturbation screens by using the mean of all perturbations as the reference ^26^ and so also removes responses shared by active perturbations; here, the reference is set by negative controls, and the shared component reflects the plate design or the reagents rather than a response to any particular cytokine.

Rhaister reproduced its published score of 0.465, slightly below the additive baseline (0.481); after correction its score fell to 0.326 (0.308 with the strict inert set) and that of the additive baseline to 0.367 (0.349), so its rank relative to the baseline did not change (Figure 5B). For State,^5^ we used the predictions released by its authors for four splits in which a cell type is held out. Under the published evaluation, State scored below the context mean in all three evaluable splits and below the additive baseline in all 12 donors in each of them (Wilcoxon *p* = 0.0005 for each cell type). After removal of the shared component, State’s score increased in the three well-sampled splits, whereas the other scores decreased. State then ranked first for CD14 monocytes, outperforming the additive baseline in 11 of 12 donors (+0.077, *p* = 0.001; +0.055 and *p* = 0.001 with the strict set), and first for naive B cells (+0.035, *p* = 0.005, 10 of 12 donors with the full inert set; ahead but not significantly with the strict set, +0.016, *p* = 0.20). For intermediate and memory B cells, State ranked second, and its gap to the baseline narrowed approximately fivefold (Figures 5C and 5D). Plasmablasts had too few donors for evaluation.

This evaluation does not penalize an error in the shared component, and State made such an error: its mean prediction for the inert proteins differed from the observed mean by 2.8 to 3.9 times the size of that mean, compared with 0.9 to 1.1 for the additive baseline. When the observed rather than the predicted inert mean was subtracted from the predictions, State fell below the additive baseline in every cell type (by 0.12 to 0.21 across both control sets; ahead in at most 1 of 12 donors), but this variant again rewarded the common component: the context mean, which carries no cytokine information, scored 0.27 to 0.44, at least as high as State (0.26 to 0.34). State therefore predicted the differences between cytokines better than the additive baseline in two cell types, but predicted the shared component, the mean response to the inert proteins in a held-out cell type, less well. No method predicted this component well: the error of the additive baseline (0.9 to 1.1) was close to that of a prediction of zero (1.0), and that of State was larger. No single score therefore captures both parts (Table S2).

### A baseline interferon effect seen in one dataset does not replicate

We finally applied the full procedure to a donor difference that looked biological, to see how far corrections within a single dataset can go. Four Parse donors have high baseline expression of interferon-stimulated genes. Among the other eight donors, the structure of the cytokine response transferred well between donors (0.67 to 0.75 of the gap between a shuffled floor and a split-half ceiling), whereas among the four interferon donors transfer was weaker (0.50 to 0.55). In B and T cells, pairs of non-interferon donors differed mainly in response size, whereas pairs of one interferon and one non-interferon donor differed more strongly in shape (Figure 6A). This persisted with the corrected pipeline (shape difference 0.07 to 0.13 between non-interferon donors versus 0.39 to 0.53 between interferon and non-interferon donors; exact permutation *p* = 0.016 to 0.040 in B, CD4 T and CD8 T cells), although not after correction for six cell types. Donors with high baseline interferon also mounted smaller responses, 0.58 to 0.73 times the size of those of the other donors across cell types.

**Figure 6.**
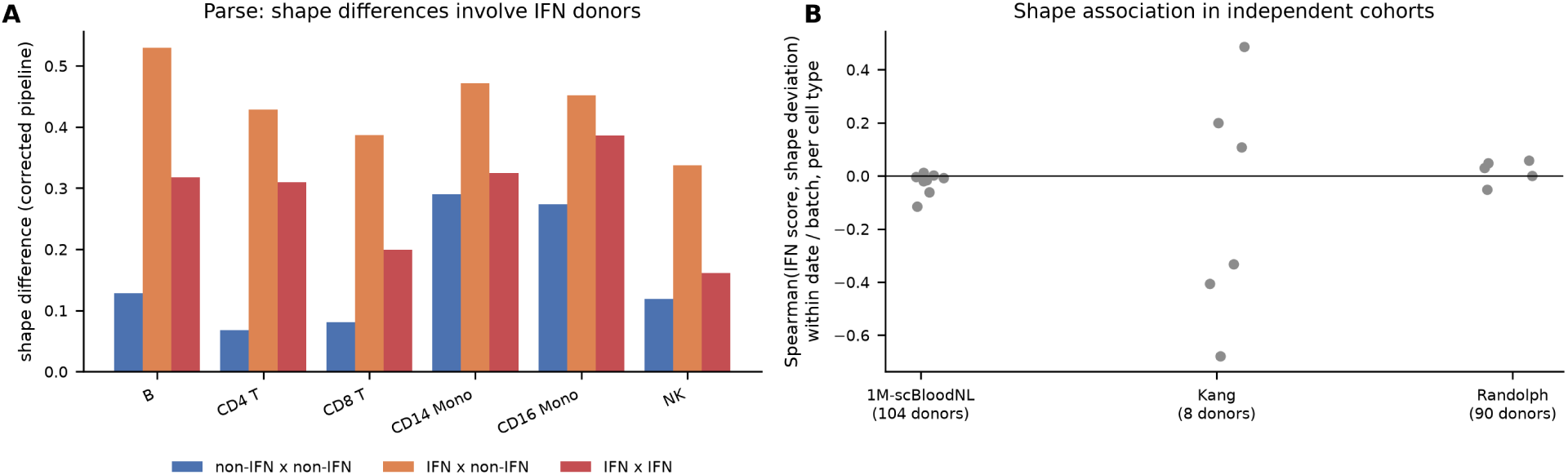
A baseline interferon effect seen in one dataset does not replicate. (A) Parse, corrected pipeline: shape difference between pairs of donors, by pair type, in six cell types. Exact permutation over all 495 possible groups of four donors. (B) Spearman correlation between the baseline interferon score and shape deviation within processing date or batch in three independent cohorts. One point per cell type; for 1M-scBloodNL, one point per cell type and chemistry with enough donors (v2, 72 donors; v3, 32 donors). In Kang et al., only CD4 T cells and CD14 monocytes had more than 19 cells per half; the other points rest on very few cells. For the fourth cohort, Aquino et al., see Figure S2C.

The shape pattern did not replicate in independent data. Baseline interferon was not associated with shape deviation in 1M-scBloodNL (104 donors with usable data) ^21^ or in 90 donors from Randolph et al.; ^22^ eight patients with lupus from Kang et al., ^27^ with very few cells per half, were uninformative. The Randolph test was well powered in CD4 T cells, with 172 cells per half, and gave a within-batch correlation of 0.05 (*p* = 0.65; Figure 6B). In the largest cohort, 222 donors from Aquino et al. stimulated for 6 h with influenza A virus (IAV) or SARS-CoV-2^20^ (donors compared within run; STAR Methods), baseline interferon did correlate with shape deviation in T cells (within-run Spearman 0.16 to 0.22, permutation *p* = 0.004 to 0.033, in CD4 T cells for both viruses and in CD8 T cells for SARS-CoV-2) and, more weakly, in myeloid cells (0.17, *p* = 0.056 for IAV; Figure S2C), and adjustment for post-thaw mortality, cytomegalovirus status and ancestry did not remove this association. An empirical null that kept each donor’s response size and noise but no difference in shape gave correlations close to zero (mean *−*0.03 to 0.04), so the association did not arise from the estimator. It did, however, depend on interferon-linked genes and on response size: in T cells it weakened and lost significance when these genes were removed (Spearman 0.03 to 0.09), and in no lineage was it significant once response size was taken into account (Figure S2C; Table S3). In myeloid cells, it remained at 0.13 without interferon-linked genes but was not significant (*p* = 0.11 to 0.14), as with all genes. Donors with high baseline interferon had much smaller responses to IAV (within-run Spearman *−*0.49 to *−*0.63), and because the response to the virus is dominated by interferon-stimulated genes, which are already elevated in these donors, a smaller response can appear as a change in shape. We therefore regard the shape result as specific to Parse until shown otherwise.

Because donors with high baseline interferon also had smaller responses, we asked whether this difference in size replicates. After removal of interferon-linked genes (STAR Methods), a smaller response persisted in Parse in CD16 monocytes, NK cells and CD8 T cells (0.70 to 0.77 times the size of the other donors’ responses with 40% of genes removed; 0.75 in CD14 monocytes, not significant) and in Aquino et al. for IAV in myeloid cells (within-run Spearman *−*0.47, similar for monocytes alone), NK cells (*−*0.19 to *−*0.22) and, more weakly, B cells, but not for SARS-CoV-2 and not in 90 donors from Randolph et al. stimulated with influenza virus (*−*0.02 to 0.14 across fractions of removed genes, except for a positive association in B cells, 0.30, in the opposite direction; Figures S2D and S7). Agreement between two datasets therefore did not amount to replication, and we do not regard this effect as established.

Independently of these results, cytokine responses shared a structure that transferred to held-out donors, filling approximately two thirds of the gap between floor and ceiling, and donors differed consistently in where each cytokine lies on that structure across three different decompositions (Figure S8).

## DISCUSSION

Three common design choices make it harder than it appears to study differences between donors and to evaluate predictive models. When each donor–perturbation combination occupies a single well shared by all cell types, well effects appear as coordination across cell types; in two independent PBMC screens, they account for two thirds to four fifths of the difference between replicate control wells in abundant cell types, and in a screen of pooled cancer cell lines for about two fifths. When control wells are grouped together, as in Parse, every comparison with the control shares a common offset, which may come from the position of the controls, from control wells being noisier than other wells or from a reagent component common to all protein wells. When the layout is identical for every donor, each perturbation carries the signature of its well into every donor, where it passes any test that uses donors as replicates. Table 1 links each feature, and the analysis choices that add to it, to the technical effects we measured, the published analyses they affect and the remedies we propose.

None of these features is specific to the datasets studied here: they follow from the plate layout, so any plate-based screen that shares this layout is exposed to the same effects, regardless of the cells or perturbations involved. We therefore view our results less as corrections to individual studies than as a general procedure: measure the technical baseline with negative controls and replicate wells, compare donors with estimators and nulls that respect the well structure, and evaluate models only on what differs between perturbations. Negative controls are the basis of established methods for removing unwanted variation, in the form of control genes and replicate samples; ^28–30^ here, proteins without receptors and identical control wells play that role for wells.

This procedure changes the interpretation of several published observations. The within-donor coordination of cytokine responses across cell types ^4^ is largely reproduced by inert proteins and by a comparison without any stimulus, because shuffling donors removes precisely what the cell types of one well share. Compound responses in the OP3 benchmark, ^3^ and its published targets, carry the row effect of each compound’s well, although the advantage this gives a position-only predictor in the benchmark metric is small; in Tahoe-100M, row effects likewise agreed between plates that carry different compounds. Wells without any active protein, including pairs of identical PBS wells and the two DMSO wells of a large cell-line screen, produce several hundred DEGs, most with small effects, so DEG counts should be interpreted relative to this floor rather than to zero, or computed with a stricter fold-change threshold, and compared only between analyses of similar power; the other effects of shared wells do not depend on DEG thresholds. None of these findings implies that the biology reported in these studies is absent; rather, the tests used could not separate it from the design.

For model evaluation, the part common to all perturbations lets a predictor that ignores the perturbation reach about 60% of the score of a simple baseline, a split-half ceiling counts reproducible well noise as signal, and a simple baseline can exceed that ceiling. Removing the shared component changed the conclusion about State: under the standard evaluation, it ranked below a predictor with no cytokine information, whereas, after correction, it ranked above the additive baseline for two of three cell types, significantly for one of them with the strict control set. Its advantage lay in the differences between cytokines; it predicted the shared component less well than the baseline, although no method predicted that component well, so the two parts are best reported separately. Several recent studies have concluded that simple baselines match or outperform complex models, both on these data and on genetic perturbation screens.^6,9,31–35^ Such conclusions should be re-examined with metrics that reward only perturbation-specific prediction, ^26^ with references derived from negative controls where these are available, and with ceilings estimated from independent wells.

For studies of inter-individual variation, the interferon result is a cautionary example: a shape difference that survived the corrections available within Parse did not replicate in the three cohorts with enough cells per donor, an apparent replication was explained by interferon-linked genes and smaller responses, and the interferon score tracked processing date, batch or run in three cohorts, more closely than ancestry in the two with ancestry information. A smaller response of high-interferon donors in myeloid and NK cells, which persisted after removal of interferon-linked genes, appeared in two datasets but not in a third: agreement between two datasets is not yet replication. A cleaner design would settle these questions more reliably than further analysis of the same data. To return to the question that motivated this work, cytokine responses shared a structure that transferred between donors, and donors differed reproducibly in where each cytokine lies on that structure (Figure S8); whether these differences are biological rather than technical can only be established with a design that separates donors from wells.

These findings lead to practical recommendations. Perturbation positions should be randomized across donor plates, so that no perturbation is permanently tied to one well and barcode; assigning positions to donors rather than to perturbations achieves the same, as in Aquino et al., where each of the 16 sample positions of a run held a donor’s NS, IAV and SARS-CoV-2 samples equally often across runs. In simulation, randomization removed the agreement of inert responses between donors, the DEGs between identical control wells and the score of a predictor that ignores the perturbation, but not the coupling of cell types within a well (Figure S4), which replicate wells and estimators built from independent halves address. Control wells should be replicated and distributed across the plate rather than confined to a single row, as recommended for high-throughput screens;^16^ in Parse, differences between control wells grew with their distance along the row, although this rests on only six wells. Proteins whose receptors are absent from the cells under study provide useful negative controls. Plate, well and processing batch should be recorded for every cell. When donors are compared, cell numbers should be equalized, noise removed with estimators built from independent halves, and differences in response shape checked against interferon-linked genes and differences in response size. Prediction models should be evaluated after removal of the component that negative controls also show, with ceilings computed from separate wells, and the accuracy of the predicted shared component should be reported separately. The donorvar package implements these analysis steps.

### Limitations of the study

Our analyses rely on the assumption that the selected proteins are inert. We restricted the strict set to proteins whose ligand-binding receptors are not detected in the main PBMC cell types, but weak effects through unmeasured receptors or in small subpopulations cannot be excluded: MPL and EPOR, the receptors for TPO and EPO, are expressed in the rare hematopoietic progenitors present in PBMCs (about 41 and 13 counts per million, respectively, in HSPCs), which were excluded from our analyses but could act indirectly on other cells; platelets, a common contaminant of PBMC preparations, also express MPL; and Megalin (LRP2) is itself an endocytic receptor that could bind ligands in the medium rather than signal. Several main findings do not depend on this assumption: the well-to-well noise, the coordination produced by a comparison without any stimulus, the DEGs between identical PBS wells and all analyses of OP3 and Tahoe-100M rely only on replicate control wells or on the plate layout. We could not identify the source of the well-specific signatures. Identical PBS wells show that a protein is not required, but barcode, handling and position remain confounded, and along row H position cannot be separated from handling order. For wells that received a protein, a reagent effect or a weak true response may add to the signature; in particular, the offset between PBS and protein wells could reflect a carrier or buffer common to all protein preparations, whose formulation was not available to us (endotoxin is unlikely; see Results). This would not change the evaluation, which removes the component whatever its source, but would make its origin a reagent rather than the layout. Only a screen in which the position of each perturbation varies between donors could separate these sources; no such plate-based screen was available to us, and our simulation shows what such a design would remove, not which source dominates in real wells. The OP3 analyses of DEG counts had less power than those in Parse, and a modest difference between the two screens remained at matched power. Our audits approximate the original pipelines: we used pseudobulk profiles; for the coordination test, we used gene space and four other spaces, including a 150-component space learned in the same way on an independent cohort, instead of the original space learned on the Sound Life cohort used by Zahid,^4^ which was not available to us; and we used the published predictions for four of the five State splits, as none were released for the fifth. Our conclusions therefore rest on comparisons within each analysis, repeated across comparison spaces and control sets. The analyses of response size were defined after the shape result had failed to replicate, and the effect was observed neither for SARS-CoV-2 nor in Randolph et al.; in Aquino et al. we analyzed the five lineages of the released data and, separately, monocytes identified by the released cell-type labels. Finally, most analyses rely on a single cytokine screen with 12 donors. The well-level findings were confirmed in two further screens, a PBMC compound screen with three donors and a cell-line compound screen (Tahoe-100M), but neither contains donor variation of the kind studied here, so the donor-level analyses rest on Parse and the four cohorts. In Tahoe-100M, the well and its position cannot be separated either, positions refer to the barcoding plate, and the row effect rests mainly on one pair of plates (see Results).

## RESOURCE AVAILABILITY

### Lead contact

Requests for further information and resources should be directed to and will be fulfilled by the lead contact, Ghazal Ghajari.

### Materials availability

This study did not generate new unique reagents.

## Data and code availability

- This paper analyzes existing data. All datasets are publicly available except that of Aquino et al., which is available on request through the Owey platform of Institut Pasteur. Accession numbers and sources are listed in the key resources table.
- All original code has been deposited at GitHub (https://github.com/Ghazal-Ghajari/p late-perturbation-confounders, release v1.0.0) and will be archived at Zenodo with a DOI before publication. ^36^
- Any additional information required to reanalyze the data reported in this paper is available from the lead contact upon request.

## ACKNOWLEDGMENTS

We thank the authors of the Parse, OP3, 1M-scBloodNL, Kang et al., Randolph et al., Aquino et al., Tahoe-100M, State and Rhaister resources for making their data, code and predictions available, and Yann Aquino for granting access to the POPCELL data and for help with the sample metadata. This research did not receive any specific grant from funding agencies in the public, commercial or not-for-profit sectors.

## AUTHOR CONTRIBUTIONS

G.G.: conceptualization, methodology, software, formal analysis, investigation, data curation, visualization, writing – original draft, and writing – review and editing. K.T.: supervision and writing – review and editing. F.A.: supervision and writing – review and editing.

## DECLARATION OF GENERATIVE AI AND AI-ASSISTED TECHNOLOGIES IN THE MANUSCRIPT PREPARATION PROCESS

During the preparation of this work, the authors used Claude (Anthropic; Claude Opus 5.5) to assist with writing analysis and plotting code and with drafting and language editing of the text, given that English is not the authors’ first language. The authors conceived and designed the study, decided which analyses to run, ran all analyses, checked the results against the data, and reviewed and edited all content. All figures were generated directly from the data by the analysis code described in STAR Methods. The authors take full responsibility for the content of the published article.

## SUPPLEMENTAL INFORMATION

Document S1. Figures S1–S8 and Tables S1–S3.

## STAR METHODS

### KEY RESOURCES TABLE

| REAGENT or RESOURCE | SOURCE | IDENTIFIER |
|---|---|---|
| *Deposited data* |  |  |
| Parse 10M PBMC cytokine screen | Parse Biosciences<sup>1</sup> | https://www.parsebiosciences.com/datasets/10-million-human-pbmcs-in-a-single-experiment/ |
| OP3 compound screen | Szałata et al.<sup>3</sup> | GEO: GSE279945 |
| 1M-scBloodNL | Oelen et al.<sup>21</sup> | https://molgenis26.gcc.rug.nl/downloads/1m-scbloodnl/ |
| Kang et al. IFN-*β* PBMC data | Kang et al.<sup>27</sup> | GEO: GSE96583 |
| Randolph et al. influenza PBMC data | Randolph et al.<sup>22</sup> | GEO: GSE162632; Zenodo: 10.5281/zenodo.4273999 |
| POPCELL PBMC data (IAV, SARS-CoV-2) | Aquino et al.<sup>20</sup> | Owey (Institut Pasteur): 10.48802/owey.e4qn-9190 |
| K562 and RPE1 Perturb-seq | Replogle et al.<sup>37</sup> | Figshare+: 10.25452/figshare.plus.20029387 |
| Tahoe-100M | Zhang et al.<sup>25</sup> | HuggingFace: tahoebio/Tahoe-100M |
| HepG2 and Jurkat Perturb-seq | Nadig et al.<sup>38</sup> | GEO: GSE264667 |
| State predictions for Parse | Adduri et al.<sup>5</sup> | HuggingFace: arcinstitute/ST-SE-Parse |
| Rhaister code and Parse statistics | Svensson et al.<sup>6</sup> | HuggingFace: tahoebio/Rhaister; tahoebio/parse-de-rhaister |
| *Software and algorithms* |  |  |
| COMPASS (method) | Liang and Singh<sup>10</sup> | Reimplemented in donorvar (this paper); method described in bioRxiv: 10.64898/2026.08.03.742643 |
| Rhaister | Svensson et al.<sup>6</sup> | HuggingFace: tahoebio/Rhaister (released code, revision 75fed20; split fewshot_donor_split_0) |
| donorvar and analysis notebooks | This paper | https://github.com/Ghazal-Ghajari/plate-perturbation-confounders |
| PyDESeq2 (version 0.5.4) | Muzellec et al.<sup>24</sup> | https://github.com/owkin/PyDESeq2 |
| Python (version 3.12.11) | Python Software Foundation | https://www.python.org |
| NumPy 2.3.5, SciPy 1.17.1, pandas 3.0.2 | Open source | https://numpy.org; https://scipy.org; https://pandas.pydata.org |
| Scanpy 1.11.5, AnnData 0.13.3 | Open source | https://scanpy.readthedocs.io |

### EXPERIMENTAL MODEL AND STUDY PARTICIPANT DETAILS

This study reanalyzed published data only; no new samples were collected. The datasets comprise PBMCs from 12 healthy donors (Parse ^2^), three donors (OP3^3^), 120 donors of the Lifelines cohort (1M-scBloodNL ^21^), eight patients with systemic lupus erythematosus (Kang et al. ^27^; a disease with elevated baseline interferon signaling), 90 donors of African and European ancestry (Randolph et al. ^22^) and 222 donors of Central African, West European and East Asian ancestry (Aquino et al.^20^), together with about 50 cancer cell lines pooled in each well (Tahoe-100M^25^). Sex, age and ancestry of the donors are reported in the original publications. Sex was not analyzed as a variable, except in Aquino et al., where it explained less than 1% of the variation in the baseline interferon score. Informed consent and ethical approval were obtained by the original studies, as described in the cited publications; the data of Aquino et al. were accessed under the data-access conditions of their authors.

## METHOD DETAILS

### Data

For donor analyses of the Parse screen, we merged fine cell types into B, CD4 T, CD8 T, CD14 monocyte, CD16 monocyte and NK cells, as in the original study;^2^ for model evaluation, we retained the fine cell types used by State. OP3 raw counts were taken from the raw/X layer of the processed file. 1M-scBloodNL data from the two chemistries (72 and 32 usable donors) were analyzed separately. For Kang et al., we used batch 2 singlets from eight donors, and for Randolph et al., the cell-level object comprising 90 donors in 15 batches. For Aquino et al., we used the processed count matrix of 222 donors 6 h after mock treatment (NS) or stimulation with IAV or SARS-CoV-2, with the five lineages of the original study (B, CD4 T, CD8 T, NK and myeloid cells) and, for a separate analysis of monocytes without dendritic cells, the released cell-type labels (CD14, CD16 and infected CD14 monocytes), and the sample metadata on run, library, ancestry, post-thaw cell mortality, cytomegalovirus serostatus, age and sex. We excluded the eight libraries of run 16, in which 16 donors had been processed a second time with CITE-seq, leaving 993,319 cells. In this design, positions were assigned to donors rather than to conditions: across runs, each of the 16 sample positions held NS, IAV and SARS-CoV-2 samples equally often.

### Pseudobulks and responses

Raw counts were summed per donor, cell type, condition and random half of cells and converted to log counts per million. Unless stated otherwise, analyses used 2,000 expressed, highly variable genes per cell type; the inert-protein comparison in Figure 1 used the 2,000 genes most highly expressed in PBS cells, so that the gene set did not depend on any response. A response was defined as the difference between a condition and its reference, either PBS or the mean of the inert proteins; for an inert protein, the reference excluded the protein itself. A pseudobulk was retained if it contained at least 10 cells and at least 5 cells in each half; the inert-protein comparison, DEG counts and model evaluation required at least 50 cells, and well-noise estimates required at least 20 cells in each half of a PBS well. A donor was excluded for a given cell type when fewer than half of the cytokines of the best-covered donor passed these filters. In the corrected pipeline, cell numbers were thinned to a common number per half for each cell type (median 34 to 348 cells per half).

### Endotoxin check

To test whether the offset between PBS wells and protein wells reflects endotoxin, we computed, for each cell type, the mean over donors of the difference between the mean of the inert proteins and PBS (log counts per million), restricted to the 5,000 genes most expressed in PBS wells. We compared the mean of this offset over 20 LPS-responsive genes (IL1B, IL1A, IL6, TNF, CXCL8, CXCL2, CXCL3, CCL3, CCL4, CCL20, PTGS2, NFKBIA, TNFAIP3, IER3, SOD2, G0S2, EREG, ICAM1, IL23A and IL10; canonical targets of TLR4 and NF-*κ*B signaling in monocytes) with 2,000 random gene sets of the same size, each gene drawn from the same decile of PBS expression as the LPS-responsive gene it replaced, and report a *z* score and a one-sided empirical *p* value for higher expression in protein wells than in PBS wells, as endotoxin would produce. Cell types in which fewer than three of these genes passed the expression filter (CD4 and CD8 T cells) were not tested. The LPS-responsive genes were lower in inert-protein wells than in PBS wells, most clearly in monocytes (*z* = *−*3.8 to *−*4.4) and less so in B and NK cells (*z* = *−*1.6 to *−*2.4).

### Responsive cytokines

A cytokine was called responsive in a cell type when the size of its response against the inert reference exceeded the 95th percentile of the sizes of differences between pairs of inert proteins in the same cell type. The threshold thus reflects how much two wells without any active protein differ.

### Decompositions

We used three decompositions: COMPASS,^10^ a single shared axis given by the top singular vector (SVD-1), and a three-dimensional shared subspace (SVD-3). Transfer to a held-out donor was scaled between a floor obtained from shuffled cytokine labels and a ceiling obtained from two halves of the donor’s cells. Donor-specific gains were compared with those obtained from another donor estimated in the same way. To validate our implementation, we reproduced on the Perturb-seq data used to introduce COMPASS ^10,37,38^ the reported cross-cell-line transfer of the perturbation-specific residual (0.22 versus 0.22 published) and obtained a comparable transfer of the shared coefficient (0.71 versus 0.66).

### Size and shape estimators

The asymmetric estimator compares one half of a donor’s cells with the mean of the other donors, and the symmetric estimator compares halves on both sides. The cross-half estimator uses only products of independent halves, *|z_d_|*2 *≈ A_d_ · B_d_ − w_d_* and *z_d_ · z_e_ ≈* (*A_d_ · B_e_* + *B_d_ · A_e_*)*/*2, where *A* and *B* are the two halves and *w_d_* is the well energy, estimated from replicate control wells as half the difference between between-well and within-well squared distances. Cell numbers were equalized across donors by binomial thinning of counts to a common number of cells per half. Where processing batches were available, donors were compared only within batch.

### Simulations

We simulated pseudobulk responses for 24 donors, 20 responsive and 6 inert conditions and 500 genes, with two halves of cells per well and 40 replicates per setting. Each true response combined a shared axis, scaled by a cytokine-specific coefficient, with a cytokine-specific direction. High-interferon donors were defined as those in the top quarter of a latent interferon state, and the observed interferon score added noise to this state. Donors were either identical, differed only in response size or, for high-interferon donors, differed in response shape. Technical settings added unequal cell numbers; well noise shared by the two halves of a well; an offset in control wells along one direction shared by all donors, with a size that varied between donors; and a processing date that raised the interferon score and shifted responses. A run was counted as significant when the Spearman correlation between the interferon score and shape deviation was positive with *p <* 0.05. Parameter values are listed in Table S1. With these values, the well accounted for about 70% of the difference between replicate control wells, within the range observed in Parse and OP3 (Figure 2A). To compare layouts, a second simulation added plate positions: 32 positions in four rows of eight, holding the 20 responsive, 6 inert and 6 control conditions, with two cell types sharing each well. Each position carried a signature that was identical on every donor plate, made of a row component (norm 2, equal to the control-well offset) and a position-specific component (SD 0.1 per gene), scaled on each plate by a factor *|N* (1, 0.5)|; well noise and sampling noise were as in Table S1. In the fixed layout, every plate used the same positions, with the control wells in the last row; in the randomized layout, each plate received an independent random assignment. Responses were computed relative to the mean of the control wells. DEGs between control wells 1–3 and 4–6 were called with paired *t*-tests across donors (Benjamini–Hochberg *q <* 0.05). The predictor that ignored the cytokine was the mean inert response of the other donors, scored by Pearson correlation with each responsive condition (leaving one donor out). Coordination was the median Spearman correlation between the inert responses of the two cell types of a donor, minus the mean over five permutations of donors. Each layout was simulated 40 times.

### Coordination audit

For each cytokine and donor, we computed the Spearman correlation between the responses of every pair of cell types in a shared space of the 3,500 genes most expressed in the PBS cells of every cell type (excluding mitochondrial and ribosomal genes) and took the median over pairs and donors. The null distribution was obtained by permuting donor labels within cell types. Technical baselines were derived from inert proteins and from a stimulus-free pseudo-response between PBS wells H7 to H9 and H10 to H12. We repeated the analysis in gene space, in a PCA of donor baseline profiles (129 components), in a PCA of all responses (150 components) and in a random 150-dimensional subspace, and in an external 150-component space: a PCA of baseline (NS) donor deviation profiles from Aquino et al. (each lineage centered across donors; 1,067 donor-lineage profiles with at least 50 cells), with genes matched by symbol (3,424 of the 3,500 genes) and Parse responses projected onto its first 150 components.

### OP3 row audit

The benchmark’s differential expression statistics were computed with limma, design ∼ 0 + sm cell type + plate name in its processing code, so plate and the combination of compound and cell type were modeled but row and column were not. For each plate and cell type, the row effect was defined as the deviation of each row from the column mean, averaged over the three control compounds in columns 1 to 3. Compound responses were computed against the plate’s DMSO mean and correlated with the effect of their own row and with the effects of the other rows. For the published targets (de train and de test of the benchmark), the log fold change of each compound and cell type was correlated with the row effect of its row averaged over plates, on the 1,882 genes shared with our gene set; *p* values came from 10,000 permutations of row labels. The position-only predictor was, for each compound and cell type, the mean of the benchmark layer (clipped signed log_10_ *p*) over the other compounds in the same row; it was scored with mean row-wise Pearson correlation, as in the benchmark, and compared with 1,000 predictors built from randomly permuted row labels, which keep the size of each group.

### Benchmark re-scoring

Rhaister was run on the Parse donor split (fewshot donor split 0) with its released code (revision 75fed20). State predictions were taken from the released outputs (eval best.ckpt). We computed a single ground truth for all methods from the raw Parse counts in State’s 2,000-gene space, normalizing each cell to 10^4^ counts followed by log(1 + *x*); this ground truth agreed with State’s own real cells at a median Pearson correlation of 0.978 to 0.990. To remove the shared component, we subtracted in each context the mean observed response over the inert proteins of the test set from the observed responses, and each method’s own mean prediction for the same inert proteins from its predictions (all methods, including Rhaister and State, predict the inert proteins of the test set). A model’s predicted difference between a cytokine and an inert protein is defined by its own prediction for that protein, which is why each method is corrected with its own inert predictions. Any predictor that is identical for all cytokines therefore yields a zero vector and scores zero. As a sensitivity analysis, we also subtracted the observed rather than the predicted inert mean from each method’s predictions, and we measured the error in each method’s predicted inert mean as ∥∥*̄p*_inert_ − *̄t*_inert_∥∥/∥∥*̄t*_inert_∥∥. Our pseudobulk evaluation scored predictions against responses computed from Parse pseudobulks as described above, whereas Rhaister’s released evaluation uses the statistics released with it (tahoebio/parse-de-rhaister); the split-half ceiling was the same score obtained when the response estimated from one random half of the cells of a query context (a held-out donor and cell type) was used as the prediction for the response estimated from the other half (at least 20 cells per half). The simple predictors were an additive baseline (the mean response to the cytokine in the training contexts plus the offset of the query context, estimated from its observed panel, that is, the cytokines measured in the query context during training), the context mean (the mean response over this observed panel) and the mean response to inert proteins in the training contexts.

### DEG counts

DEGs were called with pseudobulk DESeq2 in PyDESeq2, which treats donors rather than cells as replicates, ^39–41^ with donor and condition in the design and with the thresholds of the original study (Benjamini-Hochberg adjusted *p <* 0.05, | log_2_ fold change*| >* 0.25).^23,24,42^ Genes were tested when their expression in the pooled PBS cells of the cell type was at least 10 counts per million (for example, 8,707 to 9,401 genes in B cells, CD14 monocytes and CD4 T cells). Reproducibility of inert-protein responses was measured over 200 random splits of donors into two groups of six. Similarity between inert proteins was related to plate distance with a Mantel test. For comparisons between wells without an active protein, and for five strong cytokines against pooled PBS in the same three cell types, DEGs were also counted with | log_2_ fold change| thresholds of 0.5 and 1.

### DEGs between wells without an active protein

In B cells, CD14 monocytes and CD4 T cells, we compared each pair of PBS wells (15 pairs of wells H7 to H12), each pair of receptor-negative proteins (10 pairs) and each receptor-negative protein against pooled PBS, with the same design, thresholds and gene filter. For the power analysis, the cells of each donor, condition and cell type were placed in random order and the first 100 or 300 were retained, and the analyses were repeated with all 12 donors and with two random sets of six donors. Dependence on distance was tested with Mantel tests between DEG counts and the column distance of pairs of PBS wells (all 720 permutations of the six well labels) or the plate distance of pairs of receptor-negative proteins (all 120 permutations), and robustness by recomputing the correlation after removing each PBS well in turn.

### Tahoe-100M

Cells, raw counts and sample metadata were read from the public release (HuggingFace: tahoebio/Tahoe-100M). The DMSO wells were the samples whose drug was DMSO TF (two per plate, 28 in total; the trametinib DMSO solvate was excluded). Because cells are stored in blocks sorted by sample, only blocks whose sample range included a selected well were read, identified from the statistics in the file footers. The first entry of each cell, a special token, was removed, and counts were summed per plate, well, cell line and random half of cells (2.2 million cells in the DMSO wells; about 11 million in the compound wells read for the analyses below). The first-round barcode identified one position per well (positions 95 and 96 for the DMSO wells, the last two wells of row H, on 13 plates; positions 23 and 24, B11 and B12, on plate 13). The well share was computed as for the PBS wells, for combinations of plate and cell line with at least 100 cells in each half, on the 2,000 genes most expressed in DMSO cells, after binomial thinning of the four half pseudobulks to equal total counts. DEGs between the two DMSO wells were computed per cell line with PyDESeq2, design ∼ plate + well, for cell lines with at least six plates, with the thresholds used for Parse. Shared well noise was the Spearman correlation of the difference between the two DMSO wells (log counts per million) between two cell lines of the same plate or of different plates. For the comparison with compound responses, eight compound wells per plate were chosen at random (at the dose used on that plate), and cells were read from a random 35% of the data files. The DMSO floor was half the cross-half energy of the difference between the two DMSO wells, and the response size was the cross-half energy of the compound well against the mean of the two DMSO wells, on the same genes and with at least 50 cells per half. For the row test, all wells of plates 1 to 4 were read, and the well of each sample was taken from its first-round barcode (one position per sample, 1 to 96). We assumed that positions run row by row from A1 to H12; if they do not, each “row” is a group of 12 consecutive barcode positions, and the test still measures an effect of barcode position. The response of a well was its log counts per million minus the mean of the plate’s two DMSO wells, and the row effect was the mean response of the wells of a row minus the mean of all compound wells of the plate, for rows with at least three wells with 25 cells per half. Because plates 1 to 3 carry the same compounds in the same wells, only pairs of plates without shared compounds (each of plates 1 to 3 with plate 4) were used. Row effects of two plates were compared with Pearson correlations, and the same-row mean was tested against 1,000 permutations of the row labels, as for OP3.

### OP3 positive controls

Columns 1 and 2 of every OP3 plate hold belinostat and dabrafenib and column 3 holds DMSO, each in all eight rows. Wells with fewer than 50 cells of a cell type were excluded from DEG analyses. DEGs between two wells of the same treatment were called for all 28 pairs of rows with pseudobulk DESeq2, with plate and condition in the design (plates as replicates) and genes with at least 10 counts, and between each compound and DMSO in the same row. For well energy and split-half correlations, cells were thinned to the same even number per well within each plate and cell type (wells with at least 40 cells), and the 2,000 genes most expressed in DMSO were used. Well energy was computed as in the size and shape estimators and expressed relative to DMSO on the same plate. Responses were computed against the DMSO mean of the same plate and half of cells; for DMSO, the reference excluded the wells being compared.

### Receptor expression

For each inert protein, we listed its known receptors and co-receptors and computed their mean expression in the PBS cells of each cell type. A protein was considered receptor-negative when all of its measured receptors and co-receptors were below 5 counts per million in every main cell type, or, for Megalin, when it has no known receptor. For PSPN, the ligand-binding co-receptor GFRA4 is not measured, and the classification rests on its signaling receptor RET. The receptors of HGF (MET), PRL (PRLR), VEGF (NRP1 and FLT1), IGF-1 and Decorin (IGF1R) were expressed in at least one main cell type; those of TPO (MPL), PSPN (RET), EPO (EPOR) and EGF (EGFR) were essentially absent; and Megalin (LRP2) is itself a receptor. Three proteins were classified as uncertain: Noggin binds bone morphogenetic proteins rather than a receptor, although BMP receptors are expressed; GDNF lacks detectable GFRA1, but its alternative co-receptor NCAM1 (CD56) is highly expressed on NK cells; and FGFR1, a receptor for FGF-*β*, is expressed at low levels in the main cell types (up to 9 counts per million, in CD8 T cells) and at about 15 in MAIT cells and regulatory T cells.

### Replication cohorts

In Parse, the four donors with high baseline interferon (Donors 1, 3, 4 and 10) were those whose score, averaged over cell types, was clearly separated from the others (0.87 to 1.42 versus 0.33 or lower). The baseline interferon score was defined as the mean z-scored expression of 20 core interferon-stimulated genes in unstimulated cells: IFI27, IFI44L, IFI44, IFI6, IFIT1, IFIT2, IFIT3, ISG15, MX1, MX2, OAS1, OAS2, OAS3, RSAD2, HERC5, XAF1, EPSTI1, CMPK2, STAT1 and IFITM3. Cohorts were analyzed with the corrected pipeline, with donors compared within processing date or batch and, for Randolph et al., within ancestry. In Aquino et al., counts were summed per donor, condition, lineage and library. Each sample had been processed on two libraries, which served as independent halves (where a sample had more than two libraries, they were assigned alternately). Halves were thinned to a common number of cells per lineage (the 10th percentile over donors, at least 20), responses were computed against the donor’s NS sample on the 2,000 most variable genes with a mean of at least 10 counts per million, and each donor was compared with the mean of the other donors of the same run using the cross-half estimator. Associations were tested with Spearman correlations of ranks centered within run and two-sided tests with 10,000 permutations of the interferon score within run; partial correlations were additionally adjusted for post-thaw mortality, cytomegalovirus status and ancestry. Variation in the interferon score was partitioned with a linear model of run, ancestry, mortality, cytomegalovirus status, age and sex; the unique contribution of each factor was the increase in *R*^2^ when it was added to the other factors and was tested by permuting it. The library share was computed for NS samples as the well share for PBS wells, from four quarters of cells (two libraries times two random halves) thinned to the same number of cells.

### Interferon-linked genes and response size

Interferon-linked genes were the 20 interferon-stimulated genes listed above together with the 25%, 40% or 55% of candidate genes whose baseline expression (PBS in Parse, NS in Aquino et al.) correlated most strongly with the donor’s interferon score; the same fractions were used in both datasets. Response size was the square root of the cross-half energy relative to the mean of the other donors, per cytokine in Parse (median over responsive cytokines), per run in Aquino et al. and per batch in Randolph et al., where the two halves were random halves of cells thinned to a common number as in the corrected pipeline. In Parse, donors with high baseline interferon were compared with the others by exact permutation over all 495 groups of four donors (one-sided); in Aquino et al. and Randolph et al., size was related to the continuous interferon score within run or batch (Randolph et al.: 5,000 permutations). Size-adjusted shape tests used donor shape deviations after regression on log response size. As an empirical null for the shape estimator, each donor’s response halves in Aquino et al. were replaced by the run mean scaled to the donor’s estimated response size plus the donor’s own half-difference noise and that of another donor, so that donors differed in size but not in shape (20 replicates).

## QUANTIFICATION AND STATISTICAL ANALYSIS

Tests and the numbers of donors, cells and permutations are given in the text and figure legends. Permutation tests used 500 permutations of donor labels (coordination), 1,000 permutations of row labels (OP3 compound responses and Tahoe-100M; 10,000 for the published OP3 targets) or all 495 possible groups of four donors (interferon contrasts in Parse), 10,000 permutations of the interferon score within run in Aquino et al. and 5,000 within batch in Randolph et al. Paired comparisons between State and the additive baseline used Wilcoxon signed-rank tests on donor means. Simulations used 40 replicates per setting. Analyses were run in Python with the donorvar package and the notebooks in the code repository. Parts of the analysis and plotting code were written with the assistance of a generative AI tool (Claude, Anthropic; Claude Opus 5.5); all code was reviewed, run and checked by the authors, and every analysis and figure can be reproduced from the notebooks.

## SUPPLEMENTAL FIGURES AND TABLES

**Figure S1.**
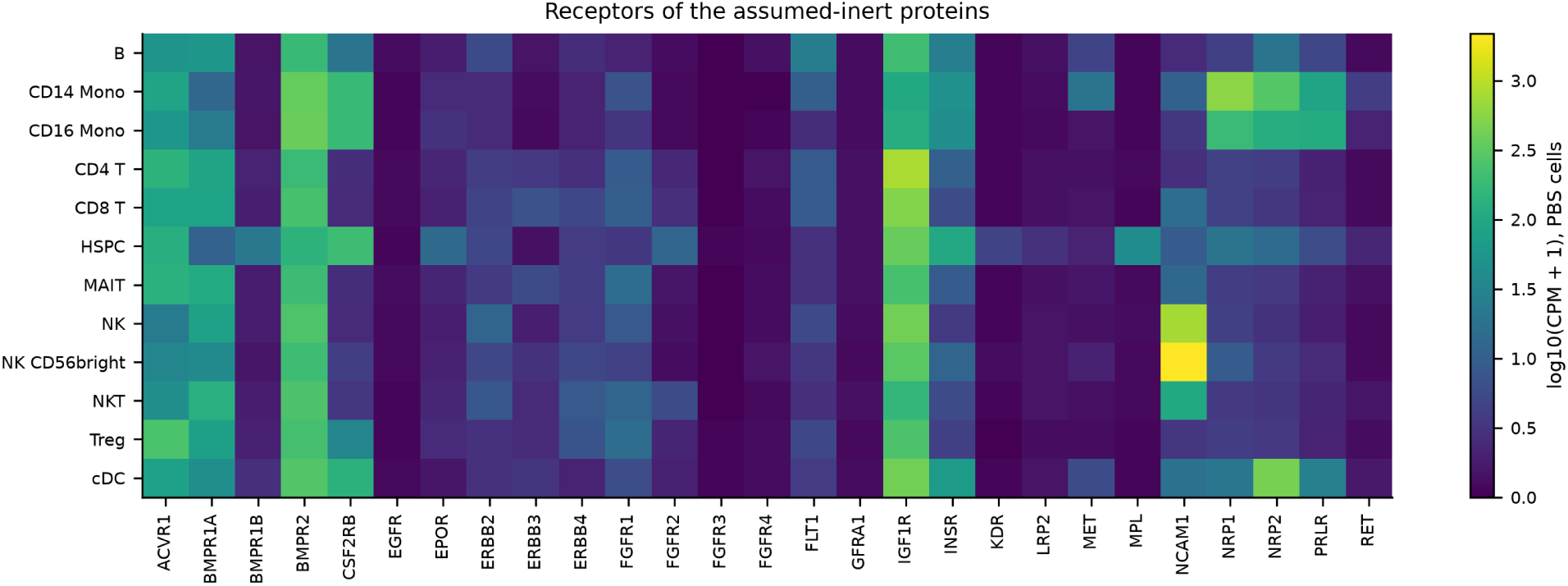
Receptors of the assumed-inert proteins, related to Figure 1. Mean expression (log_10_ counts per million + 1) of known receptors and co-receptors in the PBS cells of each cell type. Ligand-binding receptors are essentially absent for TPO (MPL), PSPN (RET; GFRA4 is not measured in this dataset), EPO (EPOR) and EGF (EGFR); Megalin (LRP2) is itself a receptor rather than a signaling ligand.

**Figure S2.**
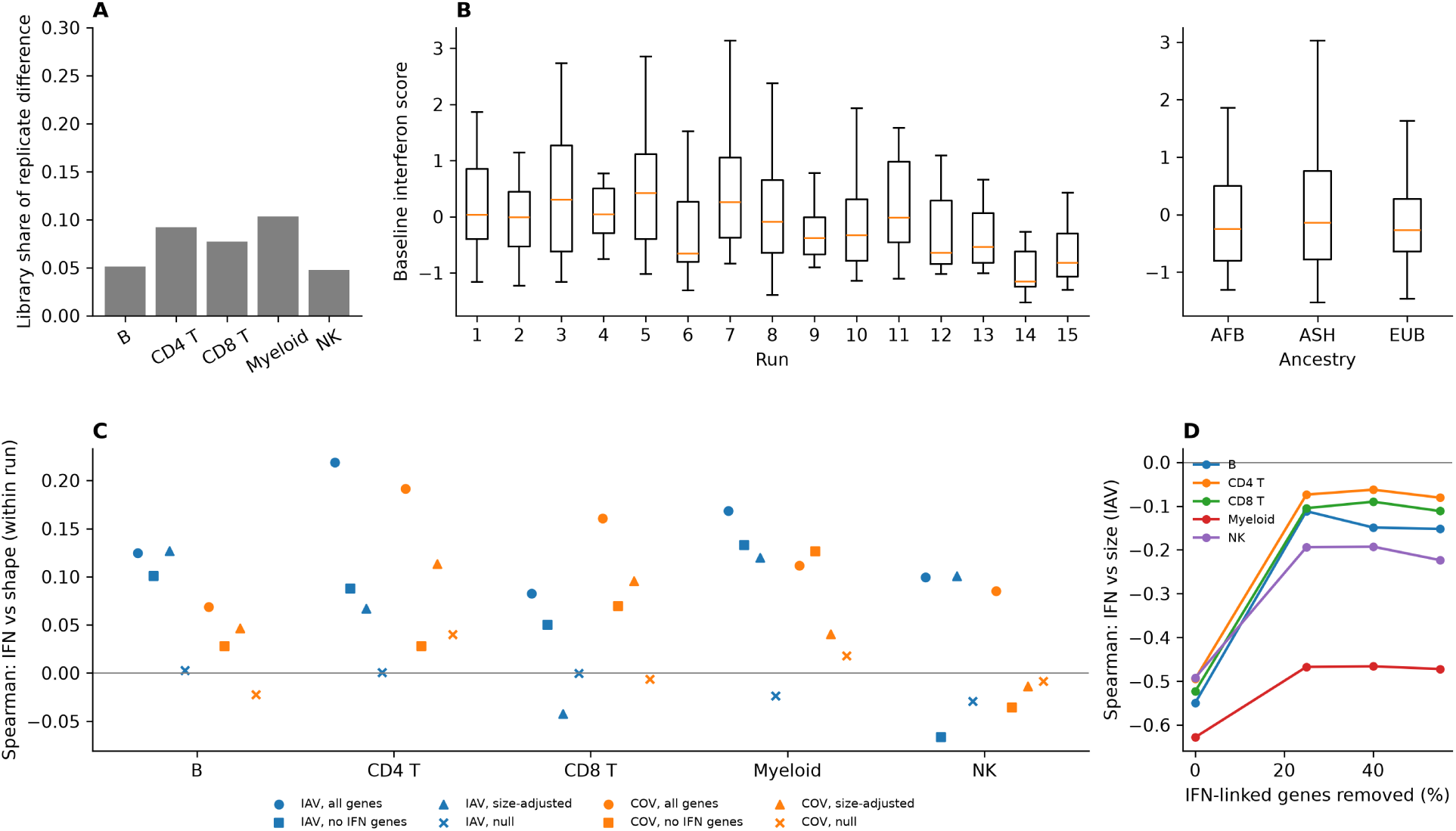
Baseline interferon, library noise and robustness of the interferon analyses in Aquino et al., related to Figures 2, 3 and 6. (A) Share of the difference between the two replicate libraries of unstimulated samples that is due to the library, (between *−* within)/between, per lineage (median over donors). (B) Baseline interferon score of 222 donors by processing run and by ancestry (AFB, Central African; EUB, West European; ASH, East Asian). Boxes, interquartile range; lines, medians. (C) Within-run Spearman correlation between baseline interferon and shape deviation for IAV and SARS-CoV-2, with all genes, without interferon-linked genes (40%), after adjustment for response size, and under an empirical null without shape differences (mean of 20 replicates). (D) Within-run Spearman correlation between baseline interferon and response size with 0%, 25%, 40% and 55% of genes removed as interferon-linked.

**Figure S3.**
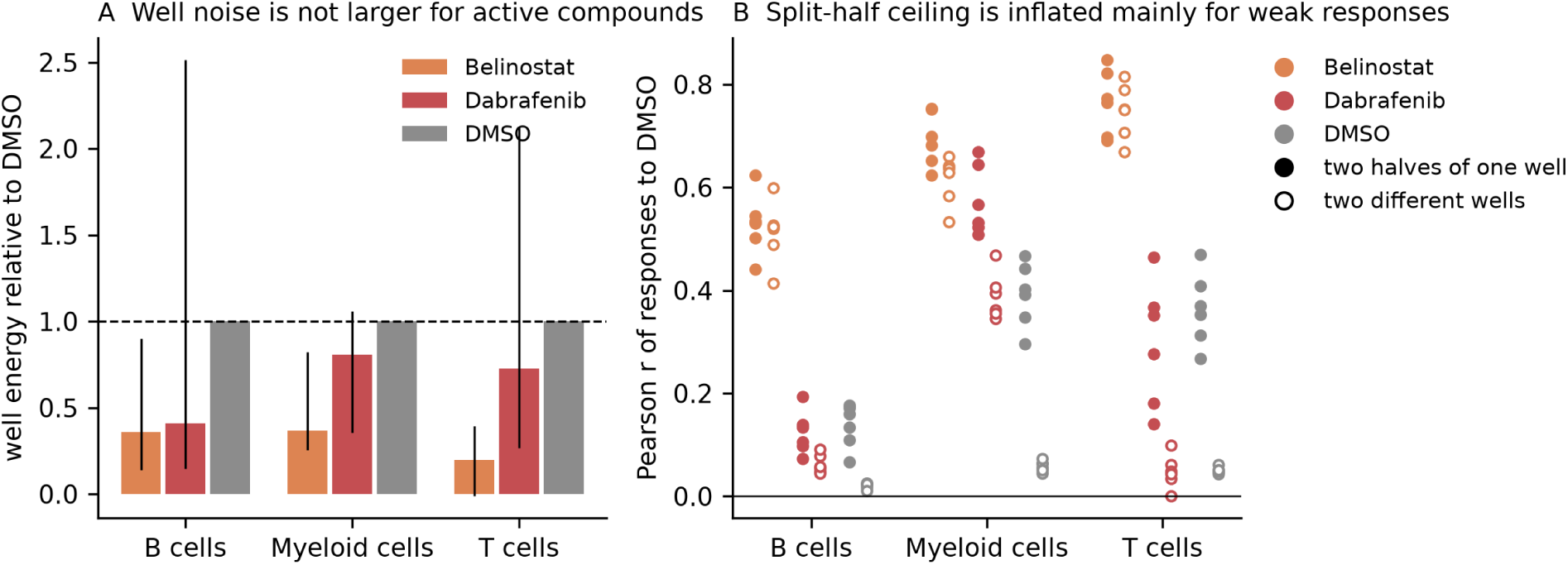
Well noise and split-half ceilings for active compounds in OP3, related to Figures 2 and 5. (A) Well energy of belinostat, dabrafenib and DMSO wells relative to DMSO on the same plate (bars, median over plates; lines, range over plates). Cells thinned to an equal number per well. (B) Pearson correlation of responses against DMSO between two halves of the same well (filled) and between two different wells of the same treatment (open). One point per plate.

**Figure S4.**
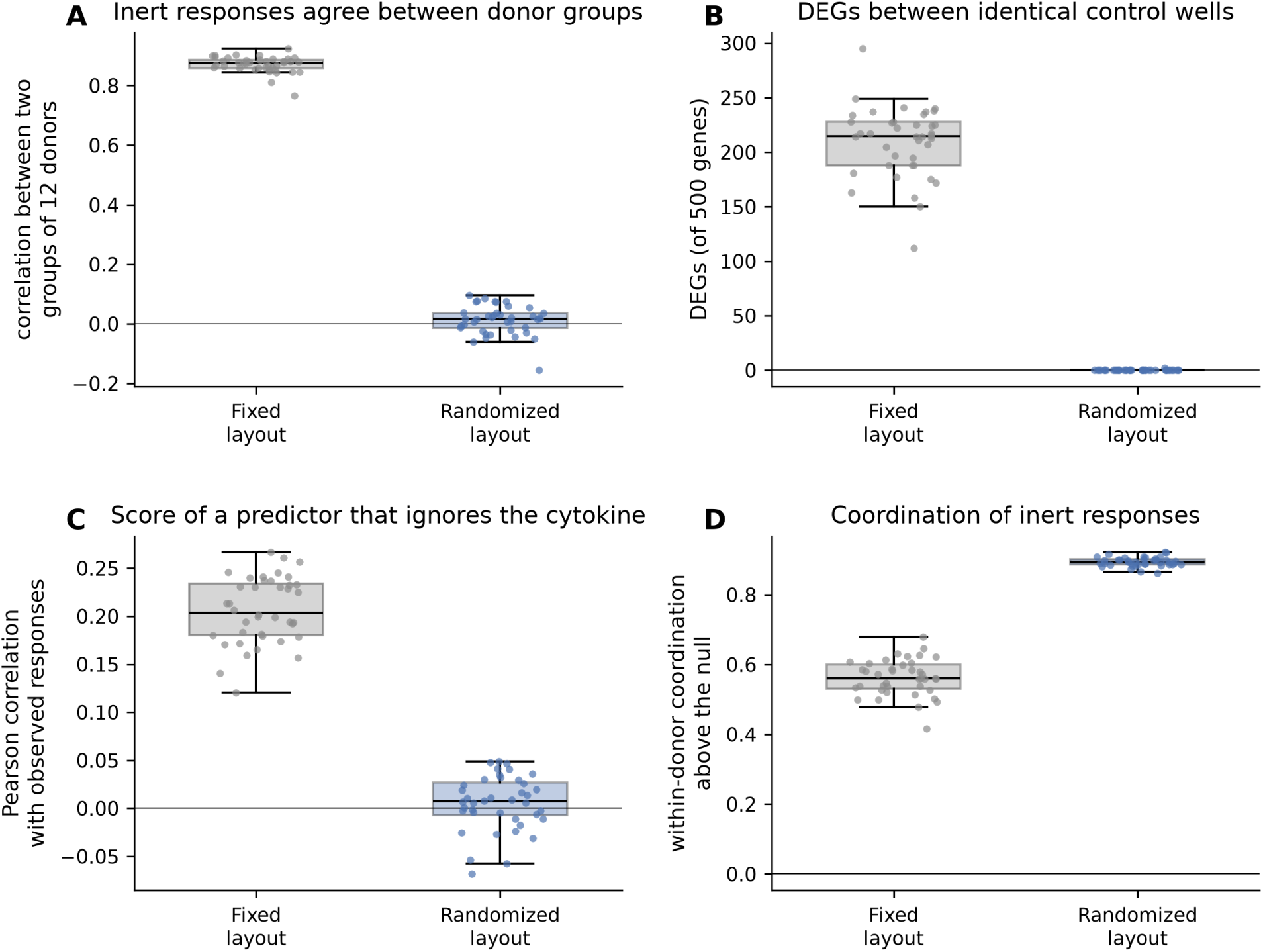
A randomized layout removes artifacts that recur across donors but not those within a well, related to Figure 3. Simulated screens of 24 donors in which each plate position carries a signature identical on every plate; fixed layout (same positions on every plate, controls in one row) versus a random layout per donor plate. 40 simulated screens per layout. (A) Mean correlation between the inert-protein responses of two groups of 12 donors. (B) Number of DEGs (of 500 genes) between two groups of three identical control wells; paired *t*-test across donors, Benjamini–Hochberg *q <* 0.05. (C) Pearson correlation between observed responses and a predictor that ignores the cytokine (mean inert response of the other donors). (D) Median within-donor Spearman correlation between the inert responses of two cell types, minus a donor-permutation null. Points, simulated screens; boxes, interquartile range; lines, medians; thin horizontal line, zero.

**Figure S5.**
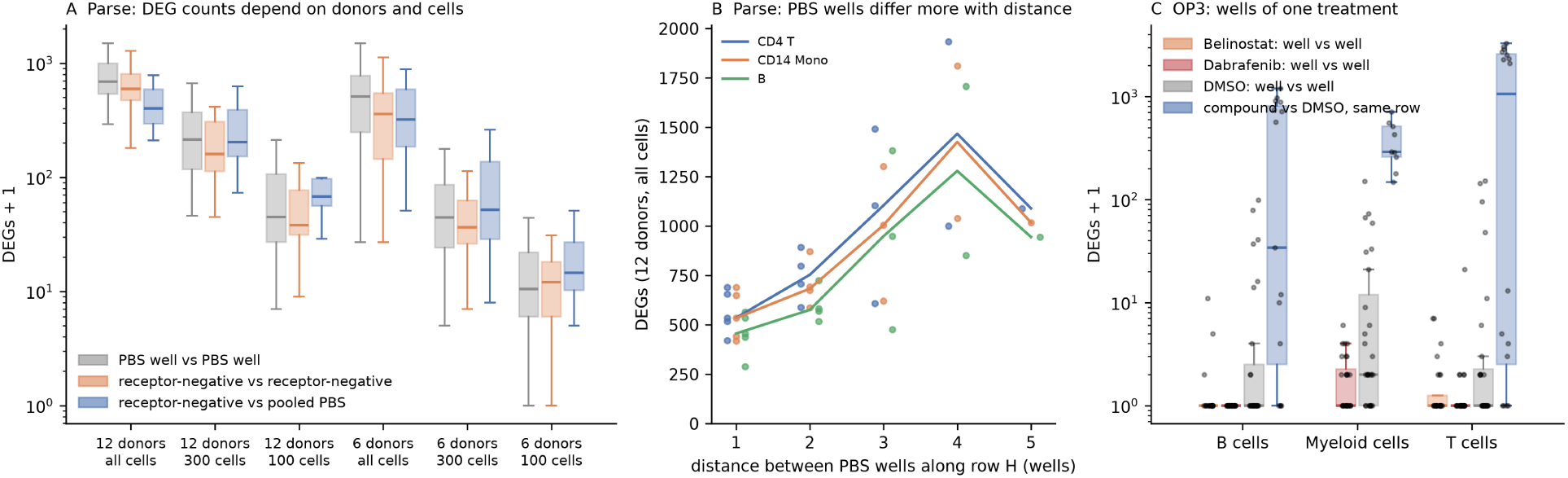
DEG counts depend on statistical power and on distance along the control row, related to Figure 4. (A) Parse: DEGs for the three comparisons of Figure 4E with all 12 donors or two random sets of six donors, and with all, 300 or 100 cells per well. Boxes, interquartile range. (B) Parse: DEGs between pairs of PBS wells against their distance along row H, with all 12 donors and all cells. Lines, medians per distance; the largest distance (5) comprises a single pair of wells (H7 and H12). (C) OP3: DEGs between two wells of the same treatment (belinostat, dabrafenib and DMSO; 28 pairs of rows) and between each compound and DMSO in the same row, with plates as replicates. Points, individual comparisons. Most belinostat wells had fewer than 50 myeloid cells and were excluded from this DEG analysis; the well-energy and split-half analyses of Figure S3 required only 40 cells per well.

**Figure S6.**
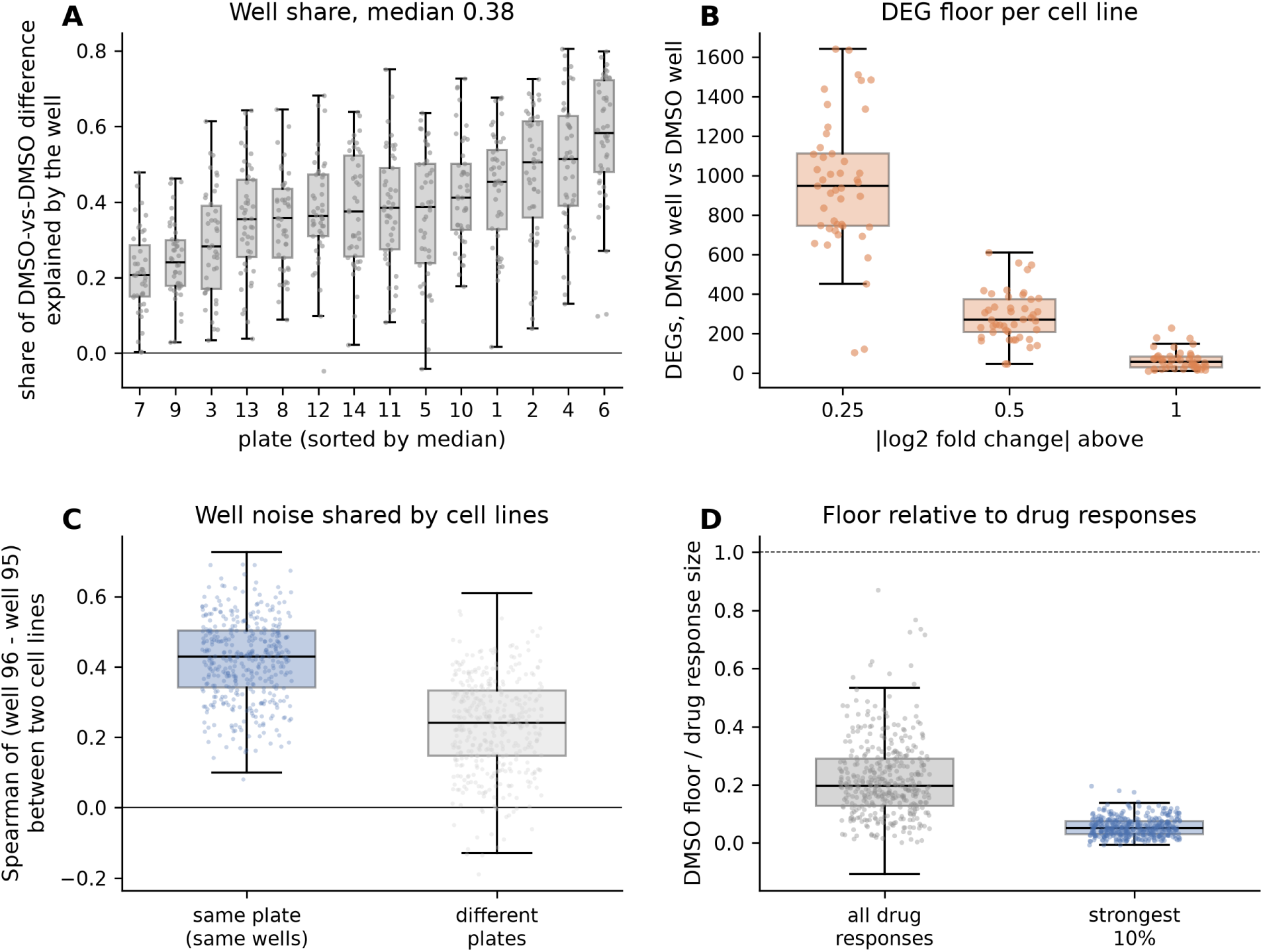
The well effects recur in Tahoe-100M, a compound screen of pooled cancer cell lines, related to Figures 2 and 4. (A) Share of the difference between the two DMSO wells of a plate that is due to the well, (between *—* within)/between, on the 2,000 genes most expressed in DMSO cells. One point per cell line; plates sorted by median. (B) DEGs between the two DMSO wells per cell line, with the 14 plates as replicates (PyDESeq2, adjusted *p <* 0.05), at three thresholds of | log_2_ fold change|. One point per cell line. (C) Spearman correlation of the difference between the two DMSO wells between two cell lines of the same plate (blue) or of different plates (grey). Points, a random subset of pairs. (D) Difference between the two DMSO wells relative to the response to a compound (eight random compound wells per plate), both as cross-half energies, for all compound wells and cell lines and for the strongest tenth of responses. Dashed line, a floor equal to the response. Boxes, interquartile range; whiskers, 1.5 times the interquartile range.

**Figure S7.**
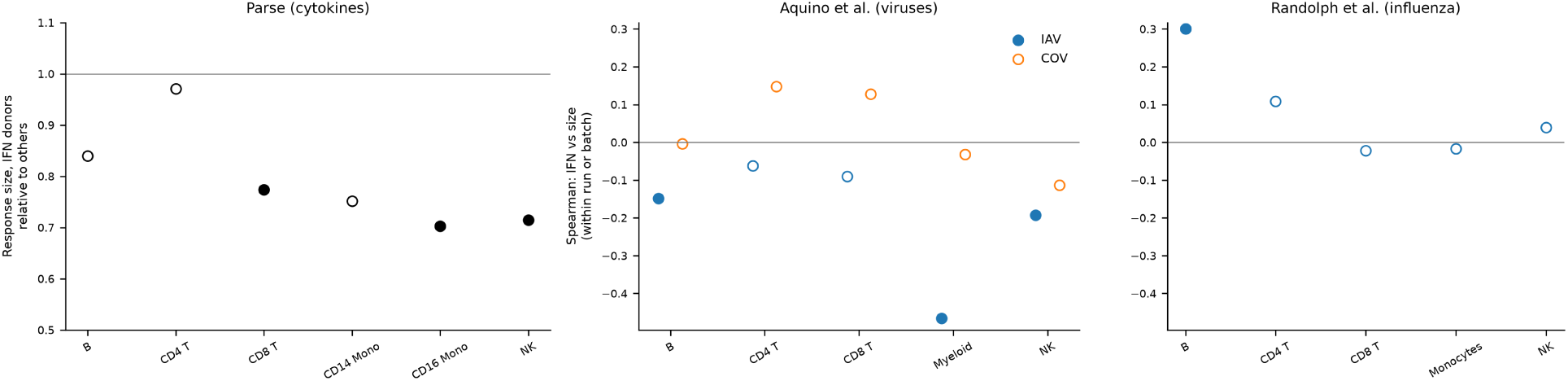
Response size of donors with high baseline interferon in three datasets, related to Figure 6. Response size after removal of interferon-linked genes (40% of genes). Parse: size relative to the other donors, one-sided exact permutation over all groups of four donors. Aquino et al.: within-run Spearman correlation between the interferon score and response size, for IAV and SARS-CoV-2. Randolph et al.: within-batch Spearman correlation, influenza A virus. Filled points, *p <* 0.05.

**Figure S8.**
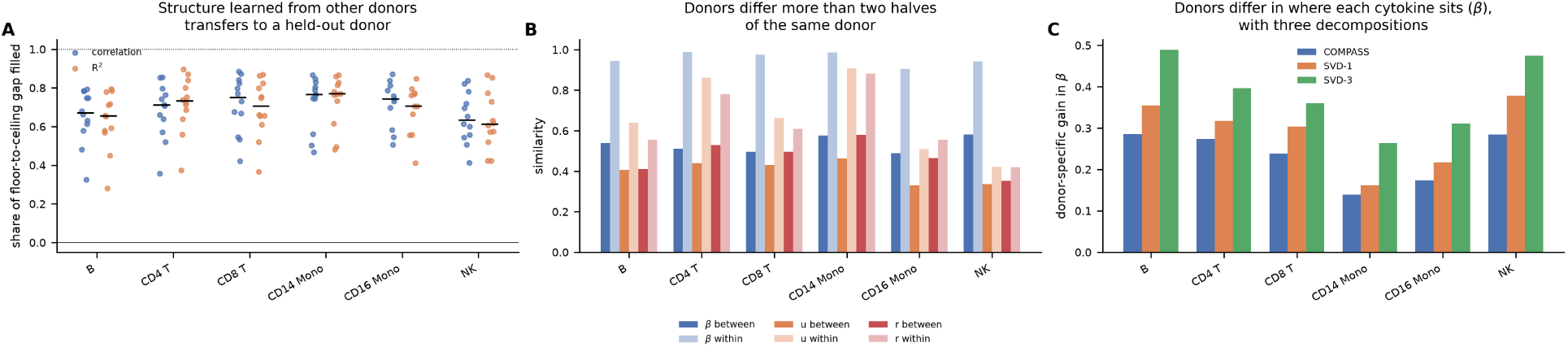
Cytokine responses share a structure that transfers between donors, related to Figure 6. (A) Leave-one-donor-out transfer of the COMPASS decomposition (shared axis, coefficients *β* and cytokine-specific residuals) learned from the other donors, scaled between a floor from shuffled cytokine labels (0) and a ceiling from two halves of the held-out donor’s cells (1), with correlation (blue) and *R*^2^ (orange). One point per held-out donor; lines, medians. (B) Similarity of the three components between donors (dark) and between two halves of the same donor (light): Spearman correlation for *β* and cosine for the shared axis *u* and the residuals *r*. (C) Gain from using the held-out donor’s own *β* over that of another donor estimated the same way, with three decompositions. Means over held-out donors.

**Table S1.** Simulation parameters.

| Parameter | Value |
|---|---|
| Donors, responsive and inert conditions, genes | 24; 20 and 6; 500 |
| Replicates per setting | 40 |
| Norm of a typical true response | 5 |
| Weight of the cytokine-specific direction | 0.5 |
| Size differences between donors (truth “size only”) | log-normal factor per cytokine, SD 0.4 |
| Shape change in high-interferon donors (truth “shape”) | added direction with norm 0.8 times the response norm |
| High-interferon donors | top quarter of the latent interferon state |
| Cells per donor | uniform, 100 to 400 |
| Cells in high-interferon donors (unequal-cell settings) | 40 |
| Sampling noise per gene | SD 1 per cell, divided by $\sqrt{\text{cells}}$ |
| Well noise (SD per gene) | 0.15 |
| Control wells | 6 |
| Control-well offset | common direction; size 0.4 times the response norm times a donor factor \|*N*(1, 0.5)\| |
| Processing date: share of donors, effect on responses, shift of interferon score | 25%; 0.4 times the response norm; 1.5 |

**Table S2.** Scores of State and two simple predictors under three evaluations, related to Figure 5. Mean pearson delta over immune test cytokines and donors for the three well-sampled held-out cell types (full inert set). Published, responses against PBS. Own inert mean, each method’s mean prediction for the inert proteins subtracted from its predictions and the observed inert mean from the observed responses (main analysis). Observed inert mean, the observed inert mean subtracted from both. Inert error, ∥∥*̄p*_inert_ − *̄t*_inert_∥∥/∥∥*̄t*_inert_∥∥. With the strict inert set, the observed-inert-mean scores were 0.29 to 0.34 for State, 0.47 to 0.55 for the additive baseline and 0.37 to 0.44 for the context mean.

| Cell type | Predictor | Published | Own inert mean | Observed inert mean | Inert error |
|---|---|---|---|---|---|
| B intermediate/memory | State | 0.287 | 0.349 | 0.301 | 2.75 |
|  | Additive | 0.509 | 0.397 | 0.477 | 0.92 |
|  | Context mean | 0.372 | 0.000 | 0.337 | 0.99 |
| B naive | State | 0.275 | 0.429 | 0.296 | 3.31 |
|  | Additive | 0.487 | 0.394 | 0.467 | 1.09 |
|  | Context mean | 0.356 | 0.000 | 0.340 | 1.19 |
| CD14 monocytes | State | 0.248 | 0.458 | 0.264 | 3.85 |
|  | Additive | 0.466 | 0.382 | 0.387 | 1.11 |
|  | Context mean | 0.349 | 0.000 | 0.265 | 1.20 |

**Table S3.** Baseline interferon and shape deviation in Aquino et al., related to Figure 6 and Figure S2C. Within-run Spearman correlation between the baseline interferon score and shape deviation, with two-sided permutation *p* values in parentheses (10,000 permutations of the interferon score within run; for the size-adjusted column, ranks were adjusted for log response size before permutation). All columns come from one run with the same thinning and permutation seeds. All genes; without interferon-linked genes (40% of genes removed); after adjustment for log response size; and the mean correlation under an empirical null without shape differences (20 replicates).

| Lineage | Virus | All genes | Without IFN-linked genes | Size-adjusted | Null (mean) |
|---|---|---|---|---|---|
| B | IAV | 0.125 (0.103) | 0.101 (0.182) | 0.127 (0.099) | 0.003 |
|  | SARS-CoV-2 | 0.068 (0.361) | 0.028 (0.713) | 0.047 (0.536) | −0.022 |
| CD4 T | IAV | 0.219 (0.004) | 0.088 (0.256) | 0.067 (0.387) | 0.001 |
|  | SARS-CoV-2 | 0.191 (0.014) | 0.028 (0.723) | 0.113 (0.151) | 0.040 |
| CD8 T | IAV | 0.083 (0.278) | 0.050 (0.507) | −0.042 (0.585) | 0.000 |
|  | SARS-CoV-2 | 0.161 (0.033) | 0.070 (0.355) | 0.096 (0.205) | −0.006 |
| Myeloid | IAV | 0.168 (0.056) | 0.133 (0.137) | 0.120 (0.179) | −0.024 |
|  | SARS-CoV-2 | 0.112 (0.156) | 0.126 (0.112) | 0.041 (0.609) | 0.018 |
| NK | IAV | 0.100 (0.197) | −0.067 (0.388) | 0.101 (0.194) | −0.029 |
|  | SARS-CoV-2 | 0.085 (0.276) | −0.036 (0.648) | −0.014 (0.862) | −0.008 |

